# Single-cell roadmap of bovine oogenesis and somatic niche interactions during fetal ovarian development

**DOI:** 10.64898/2026.08.24.745837

**Authors:** Carly Guiltinan, Ramon C. Botigelli, Rachel B. Arcanjo, Justin M. Smith, Courtney K. Grimm, Sarah K. Plummer, Brendan P. Keough, Mark N. Paulsen, Sandeep K. Rajput, Benjamin P. Beaton, Anna C. Denicol

**Author notes:** Current address: Department of Biomolecular Engineering, University of California, Santa Cruz, Santa Cruz, CA, USA. Current address: Departamento de Genética e Evolução, Universidade Federal de São Carlos, São Carlos, SP, Brasil. These authors contributed equally (Carly Guiltinan, Ramon C. Botigelli).

## Abstract

The major events of female germline establishment, from primordial germ cell (PGC) specification to assembly of primordial follicles, occur during embryonic–fetal development. This study presents a single-cell RNA-sequencing atlas of the bovine fetal ovary at four gestational timepoints: estimated day 50, and timed pregnancies at days 70, 90, and 120, capturing the progression of PGCs through commitment, meiotic entry, and early oocyte growth. Fourteen transcriptionally-distinct cell populations were identified, including stromal, epithelial, endothelial, immune, somatic support cell, and germ cell lineages. Sub-clustering of the germ cell population resolved six developmental stages (PGCs, transitioning oogonia, proliferative oogonia, committed oogonia, meiotic prophase I oogonia, and oocytes), while that of the somatic support cell compartment revealed five granulosa cell subtypes (steroidogenic, pre-granulosa 1, pre-granulosa 2, pre-granulosa 3, and epithelial cells). Trajectory analysis reconstructed the developmental path of PGCs to oocytes, with sequential activation of meiotic and oocyte-specific gene programs. Representation of all six germ cell stages at day 120 of fetal development pointed to asynchronous oogenesis in the forming ovary, which was validated and shown to be region-specific by protein immunolocalization. Intercellular signaling networks between germ cells and the somatic niche were mapped, revealing strong interactions through BMP, KIT, IGF, IGFBP, WNT, and MDK pathways with temporal specificity across gestational ages. The bovine germ cell and pre-granulosa cell subtypes demonstrate significant transcriptional parallels with similarly-staged cells from human fetal ovaries, establishing the cow as a reliable model for human germ cell and ovarian development. Collectively, these data provide a developmental roadmap for bovine oogenesis at the single-cell resolution that advances fundamental understanding of gametogenesis and informs strategies for advanced assisted reproduction.

## Introduction

Females are born with a reserve of follicles—comprised of oocytes and somatic support cells—that serve as a finite resource and determine the reproductive lifespan. Oocyte development is a highly coordinated, multi-step process that initiates during early embryogenesis and proceeds through several critical steps during fetal growth before reaching quiescence until puberty. While the key steps of germline emergence are relatively conserved among mammals, cross-species comparisons have shown important distinctions in the molecular regulation of this process. Following their specification in the peri-gastrulation epiblast via a tripartite transcription factor network, primordial germ cells (PGCs) directionally migrate through the forming gut to the genital ridges^1–5^. Upon reaching the genital ridges (coined bipotential gonads upon PGC arrival), germ cells interact with the somatic compartment, which guides their development toward competent gametes^6^. Namely, PGCs acquire expression of germline commitment markers, followed by a gradual loss of pluripotency and specification factors^7,8^. Moreover, PGCs begin proliferating rapidly in the gonad to expand upon a relatively small founder population^5,9^. Somatic sexual differentiation based on genetic sex guides ovarian formation, including the emergence of pre-granulosa cells, which support advanced development of PGCs into oogonia^10^. Incomplete cytokinesis during mitosis of oogonia leads to the formation of a connected network of germ cells surrounded by pre-granulosa cells, known as ovigerous cords, which facilitate coordinated germ cell development by enhanced cell communication^11^. Uniquely in females, retinoic acid signaling from forming granulosa cells initiates meiosis in the germline until arrest at prophase of meiosis I ^11,12^. As germ cell nests breakdown, a considerable number of oogonia transition into nurse cells and then undergo programmed cell death to regulate germ cell number and support the surviving cells^13^. The remaining oogonia proceed through oocyte development and follicle assembly, during which oocytes are individually encased by a single layer of granulosa cells to form primordial follicles. These follicles represent the reproductive reserve of a female, as germ cell proliferation ceases during fetal development. They remain quiescent until puberty, at which time follicle activation and development supports the advancement of selected oocytes into cells that are competent for fertilization.

Initial studies of bovine oogenesis traced germ cell location and developmental status by histological analysis of samples throughout embryonic and fetal growth (days 18–210 of pregnancy), providing important context about the approximate timeframe of major steps of germline establishment^14–16^. Following their specification by day 18, bovine PGCs migrate until approximately day 27–31^16^. Upon their colonization of the gonad, which is sexually indifferent until approximately day 39, germ cells have a PGC-phenotype between days 35–55 and proceed through oogonial development and peak mitotic activity between days 56–80. Meiotic activity, as well as formation of oocytes, bi/multi-nucleated germ cells, and degenerating germ cells, occur between days 80–130^15^. The first primordial follicles could be identified in the bovine fetal ovary by day 90, increasing in number until day 140, when the first primary follicles appear, regulated by the steroidogenic activity of the forming ovary. From this subset of growing primary follicles, occasional secondary follicles can be observed from day 210^14^. Within this window, others have explored the temporal expression pattern and function of specific genes in bovine germ and somatic gonadal cell development via protein immunolocalization, quantitative gene expression analysis, and gene knock-out studies^17–21^. Recent reports have harnessed RNA-sequencing technologies to uncover the molecular profile of bovine germ cells and their surrounding niche throughout key timepoints of development^2,22,23^. Our group recently demonstrated the timing and molecular regulation of bovine primordial germ cell specification by day 16 and migration onset by day 20 of embryogenesis^2^. Transcriptomic profiling of the fetal ovary at estimated day 50 of development indicated that the majority of germ cells were in early stages of development, with a subset of cells transitioning toward germline commitment and a small fraction acquiring responsiveness to meiotic cues^23^. Importantly, these studies identified commonalities between bovine and human germ cell molecular profiles, highlighting a conserved program of germline establishment from specification to initial development at the gonad^2,23^.

The potential applications of understanding the molecular mechanisms of bovine fetal germline development extend well beyond basic biology. In vitro gametogenesis (IVG), or the derivation of functional gametes from pluripotent stem cells (PSCs), has been achieved in mice through a series of landmark studies^24,25^, but this milestone has not been achieved in any non-rodent species to date^26^. An efficient protocol to generate bovine PGC-like cells from PSCs was recently described, however these cells could not progress beyond this stage in reconstituted ovaries with murine somatic gonadal cells^27^. This gap is attributable in large part to an incomplete molecular blueprint of in vivo germ cell development—without knowing the milestones to be recapitulated, empirical protocol optimization is severely constrained. For cattle, the economic stakes are substantial; in vitro breeding schemes that incorporate IVG could shorten the generational interval by an order of magnitude relative to current assisted reproductive technologies, with simulations suggesting a tenfold acceleration in genetic progress compared to conventional embryo transfer utilizing genomic selection^26,28,29^. Additionally, such an advancement would be relevant to the conservation of endangered ungulate species from which induced PSCs (iPSCs) are available^30^.

Here, we traced the major steps of oogenesis in the bovine fetal ovary by performing single-cell RNA-sequencing (scRNA-seq) from timed pregnancies at days 70, 90, and 120, which was integrated with published data from estimated day 50, with four objectives: (1) to define the single-cell transcriptomic landscape of the bovine fetal ovary across this critical developmental window; (2) to characterize the transcriptional identity and developmental trajectory of germ cells from PGC to oocyte; (3) to trace the emergence of granulosa cells leading up to follicle assembly, and (4) to map the intercellular signaling architecture linking germ cells to their somatic niche at the fetal stages. The resulting atlas provides a temporal, developmental roadmap for bovine oogenesis with single-cell resolution.

## Material and Methods

All reagents and consumables were purchased from Thermo Fisher Scientific unless otherwise specified.

### Bovine fetal ovary collection and dissociation for single-cell RNA-sequencing

Bovine embryos were produced in vitro using X-skewed semen following established protocols. On day 7, embryos were transferred into estrus-synchronized recipient cows. Pregnancies were maintained for 68–70 (n = 3), 86–90 (n = 3), and 118–120 (n = 4) days. Recipients underwent cesarean section for fetal collection. Fetal ovaries were immediately dissected and washed in ice-cold phosphate-buffered saline (PBS). Samples were transported in PBS on ice to the lab for processing, which began within a maximum of 90 min from collection. One ovary from each pair was cut in half crosswise and put into a 1.5 mL tube with 0.5 mL of dissociation solution comprised of 5 mg/mL collagenase type IV (LS004188, Worthington Biochemical Corporation) and 10% v/v 7 mg/mL DNase I (DN25, Sigma) in RPMI 1640 medium. All centrifugation steps were performed at 250 x g for 4 min. Samples were finely minced with sterile scissors. An additional 1 mL of dissociation solution was added to each tube and gently mixed with a P1000 pipette 5–10 times. Tubes were incubated in a 37 °C water bath for 20 min with a gentle mix 10–15 times by P1000 every 5 min and then centrifuged. Supernatant was removed, and the pellet was resuspended in 450 µL TrypLE Express Plus 50 µL DNase I. Tubes were incubated at 37 °C for 5 min with a mix by flicking halfway through. Then, TrypLE was inactivated by dilution with 1 mL HBSS plus 10% fetal bovine serum (FBS), and the tubes were centrifuged. Supernatant was removed, and pellets were resuspended in 500 µL 3 mg/mL Pronase (P8811, Sigma) plus 50 µL 7 mg/mL DNase I. Tubes were incubated at 37 °C for 6 min with a mix by flicking at halfway and before centrifugation. Supernatant was removed, and each pellet was resuspended with 100 µL ACK lysis buffer (A1049201, Thermo Fisher) for 40 sec, immediately diluted with 1 mL HBSS + FBS + 100 µL DNase I solution, and then centrifuged. The supernatant was discarded, and the pellets were resuspended in 1 mL HBSS + FBS. This suspension was run through a 40 µm filter pre-rinsed with HBSS + FBS into a 50 mL tube. The 1.5 mL tubes and filters were rinsed 2 times with 1 mL HBSS + FBS. The new 50 mL tubes were centrifuged, supernatant discarded, and resuspended in HBSS + FBS. That cycle was repeated for a wash, with the final resuspension in 1 mL moved back to a 1.5 mL tube. Cell number and viability were assessed in triplicate by an automated cell counter (DeNovix) diluted 1:1 with 0.4% Trypan blue solution, and the mean cell number was used for final dilution (1000 cells/µL) for 10X cell suspension.

### Single-cell RNA-sequencing, alignment, quality control, and preprocessing

Bovine ovarian suspensions were processed using the 10X Genomics Chromium Controller with a targeted cell recovery of 5,000 cells per sample. Libraries were prepared using the 10X Chromium Next GEM Single Cell 3’ kit v3.1. Sequencing was performed with Illumina NextSeq 2000 set to 28 bp (Read 1), 10 bp (Index 1 and 2), and 90 bp (Read 2) using NextSeq 1000/2000 P2 reagents (100 cycles, 20k reads per cell). Raw sequencing data from one bovine fetal ovarian sample at approximately 50 days (age of fetus estimated by crown–rump length) from Soto and Ross (2021) was included in the analysis. Alignment of sequencing reads to the bovine genome (NCBI ARS-UCD2.0 with manual SOX17 annotation from Ensembl ARS-UCD1.2), and initial analyses were performed using the CellRanger pipeline v8.0.0 (10X Genomics) ^31^, followed by the Seurat package (v5.3.0)^32^ of R. Seurat objects from each sample (4 samples) were independently created and processed according to standard Seurat protocols. Firstly, single cells with a number of detected genes (nFeature_RNA) above 800 from Soto and Ross (2021) sample and above 300 from our samples and below 6,000 were retained to exclude low-quality cells. Then, doublets or multiplets cells were identified with the scDblFinder R package and excluded^33^. After normalization of the Seurat object, we selected the 5000 most variably expressed genes using the ‘FindVariableFeatures’ command. We then used the ‘ScaleData’ and ‘SCTransform’ functions of Seurat to exclude individual heterogeneities between samples. Data integration was performed using the ‘Harmony integration’ method (Harmony, 1.2.4). Fifty dimensions were selected via the ElbowPlot method for the ‘FindNeighbors’ and ‘FindClusters’ functions. Seurat pipeline, a resolution of 0.6 and 13 dimensions were used to identify the 26 clusters. We identified the 13 major cell types, which were manually annotated by inspection of known marker gene expression and supported by differential expression analysis using the ‘FindAllMarkers’ function (Wilcoxon rank-sum test, adjusted p < 0.05, log2FC threshold = 0.25, min.pct = 0.1).

### Gene ontology (GO) and pathway enrichment

Over-representation analysis (ORA) and gene set enrichment analysis (GSEA) were performed with the SCP package against the GO Biological Process (GO:BP) database. ORA used top cluster marker genes (log2FC > 0.5, adjusted p < 0.05); GSEA used all genes ranked by log2FC. Statistical significance threshold: adjusted p < 0.05.

### Sub-clustering of germ cell and somatic support cell populations

Cells assigned to germ cell and granulosa progenitor cell lineages were separately extracted and re-clustered using the same workflow. After sorting germ cells, 3 clusters were excluded from the analysis due to a low-quality profile. This yielded six germ cell subtypes and four somatic support cell subtypes. Sub-clustering results are presented in Figures 2 and 3, respectively.

### Developmental trajectory and pseudotime analysis

Pseudotime trajectories were inferred using Slingshot^34^ after transferring Seurat objects to SingleCellExperiment format. Briefly, Slingshot constructs a minimum spanning tree across cluster centroids to define lineage trajectories, then fits simultaneous principal curves to assign pseudotime, a scalar value indicating each cell’s relative position along the developmental path. Root nodes were assigned based on biological priors: Steroidogenic cells for the pre-granulosa lineage and PGCs for the germ cell lineage. Genes with significant pseudotime-dependent expression (graph_test, q < 0.05) were visualized as smoothed curves stratified by lineage. Pseudotime-ordered heatmaps were generated with genes grouped by expression dynamics using unsupervised clustering.

### Transcription factor regulon analysis (SCENIC)

Gene regulatory networks and regulons were inferred using the SCENIC workflow in R (GENIE3, RcisTarget, and AUCell packages), applied to the germ cell subset spanning D50e to D120 of bovine fetal ovary development. Raw RNA counts were extracted from the Seurat object. Genes were retained for regulatory network inference using the geneFiltering function (SCENIC package) with two criteria: a minimum counts per gene (2 * 0.01 * number of cells; equals 62 counts in the dataset) and minimal sample (minimal of 20 cells). Co-expression networks were built with GENIE3 (500 trees per gene), and regulons were defined by linking each transcription factor to its top 5, top 10, and top 50 correlated target genes. Because a bovine-specific motif database was not available, the default human (hg19, HGNC) cisTarget ranking databases distributed with SCENIC were used as a proxy for transcription factor motif enrichment, comprising two feather-format databases based on the 7-species mc9nr motif collection: regions spanning 500 bp upstream of the transcription start site (TSS), and regions extending 10 kb upstream and downstream of the TSS. Regulon activity was scored per cell with AUCell, and cells were classified as regulon-active or -inactive using automatic binarization. Mean regulon activity was calculated across annotated cell types and developmental timepoints, and cell-type specificity of each regulon was assessed with the regulon specificity score (RSS).

### Cell–cell communication analysis

Intercellular signaling was inferred by ligand–receptor pair expression using CellChat v2^35^ on the subset dataset (by age) from the full integrated dataset. After that, a comparison was made between all-time points evaluated. Signaling pathways with biological relevance to germ cell development, such as BMP, KIT, and IGFBP, were examined in detail and visualized as network diagrams with arrow strength proportional to interaction score.

### Bovine fetal ovary collection and immunolocalization of germ cells proteins

Bovine pregnant reproductive tracts and fetuses were purchased from local abattoirs and transported to the laboratory submerged in ice within 2 to 3 h. Tracts were dissected, and fetal age was estimated based on the crown–rump length according to DesCôteaux et al. (2009). Sex of the fetuses was identified by internal and external morphology. Ovaries from fetuses between 60 and 120 days of development were fixed with 4% paraformaldehyde overnight at 4 °C. Fixed fetal ovaries were cryoprotected in 30% sucrose until tissue sank, then embedded in Tissue-Plus OCT compound (Fisher HealthCare). Cryosections (10 µm) were cut on a cryostat, mounted on Superfrost Plus slides, and stored at −20 °C. Before staining, sections were brought to room temperature, rinsed twice with TBS and once with distilled water to remove residual OCT, then subjected to antigen retrieval in Tris-based unmasking solution (pH 9.0, Vector Laboratories) for 20 min at ∼93 °C on a steamer. Once cooled, sections were washed with TBS-T [TBS + 0.025% (v/v) Triton X-100], permeabilized for 10 min with 0.5% Triton X-100 in TBS, and washed again with TBS-T. Blocking was performed for 1 h at room temperature in 10% (v/v) normal donkey serum, 1% (w/v) bovine serum albumin (BSA), and 0.3 M glycine in TBS-T. Primary antibodies were diluted in 1% BSA in TBS-T and applied overnight at 4°C in a humidified chamber. Sections were washed three times with TBS-T, then incubated with secondary antibodies in the same buffer for 1–2 h at room temperature in the dark. DNA was labeled with Hoechst 33342 (1:1000, 15 min, room temperature, dark), and slides were mounted with ProLong Gold Antifade, cured overnight, and sealed with clear nail polish. Antibodies and dilutions are listed in Table 1.

**Table 1.** Antibodies used for protein immunolocalization analysis of bovine fetal ovaries. All antibodies were reconstituted according to manufacturer protocols, when applicable.

| Primary antibody | Manufacturer | Catalog number | Dilution |
| --- | --- | --- | --- |
| Goat anti-POU5F1 | R&D Systems | AF1759 | 1:100 |
| Rabbit anti-DAZL | Santa Cruz | Sc-390929 | 1:500 |
| Secondary antibody | Manufacturer | Catalog number | Dilution |
| Donkey anti-Goat 568 | Invitrogen | A11055 | 1:500 |
| Donkey anti-Rabbit 488 | Invitrogen | A31573 | 1:500 |

### Image acquisition and processing

Stained sections were imaged on an ImageXpress Micro Confocal microscope (Molecular Devices, San Jose, CA) at 20x magnification in tiled scanning mode with 10% overlap, capturing the entire section at high resolution. Tiles were stitched into a single montage for whole-section evaluation. Exposure settings, light intensity, and brightness/contrast adjustments were established using isotype controls for each antibody in MetaXpress software relative to those controls.

## RESULTS

### Population composition of day 50–120 bovine fetal ovaries revealed at the single-cell resolution

To produce a single-cell roadmap of germline development within the forming bovine fetal ovary, we performed integrative analysis of scRNA-seq data generated from bovine ovaries at days (D) 70, 90, and 120 of fetal age, along with a published dataset from day 50 of estimated fetal age (D50e)^23^. After quality control, doublet removal, and low-quality cluster exclusion, a total of 50,070 cells were retained for downstream analysis: 13,782 from D50e; 13,287 from D70; 12,425 from D90; and 13,576 from D120 (Supplementary Figure 1A). We applied UMAP dimensionality reduction to the Harmony-integrated dataset to visualize transcriptional relationships across cells (Figure 1A–B). Unsupervised clustering and marker-guided annotation identified 13 transcriptionally-distinct cell populations (Figure 1C): stromal cells, stromal fibroblasts, stromal proliferative cells, granulosa progenitor cells, proliferative steroidogenic cells, smooth muscle cells, epithelial cells, endothelial cells, germ cells, macrophages, natural killer (NK) cells, lymphocytes, and erythroblast cells. The relative contribution of each timepoint to each cluster is shown in Figure 1D, which illustrates temporal shifts in cell type representation across the D50e– 120 window. Complete differential gene expression (DGE) (Supplementary Figure 1), canonical marker genes, including *PDGFRA*/*TCF21* for stromal populations, *DDX4*/*DAZL* for germ cells, *WNT6*/*DACH1*/*KITLG*/*CYP19A1* for granulosa progenitors, and *PECAM1*/*CDH5* for endothelial cells (Figure 1E), and gene ontology (GO) enrichment analysis (Supplementary Figure 2) were used for cell type annotation. Stromal cells, defined by high expression of *COL1A2, COL12A1, LTBP4,* and *IGFBP3*, were the most abundant somatic population across all timepoints. The granulosa progenitors cluster expressed *STAR, LHCGR*, and *CXCL14*, consistent with a transitional identity between stromal and steroidogenic lineages with theca-like identity. Germ cells, identified by canonical markers including *POU5F1, PRDM1, DDX4*, and *DAZL*, formed a distinct, transcriptionally-isolated cluster whose relative abundance changed across gestational times, reflecting the dynamics of proliferation, meiotic entry and arrest, and germ cell attrition during follicle assembly (Figure 1D). Immune cell populations (macrophages, NK cells, and B lymphocytes), endothelial, erythroblast, and smooth muscle cells were clearly separated and are consistent with expected populations within the fetal gonad vasculature and immune niche.

**Figure 1.**
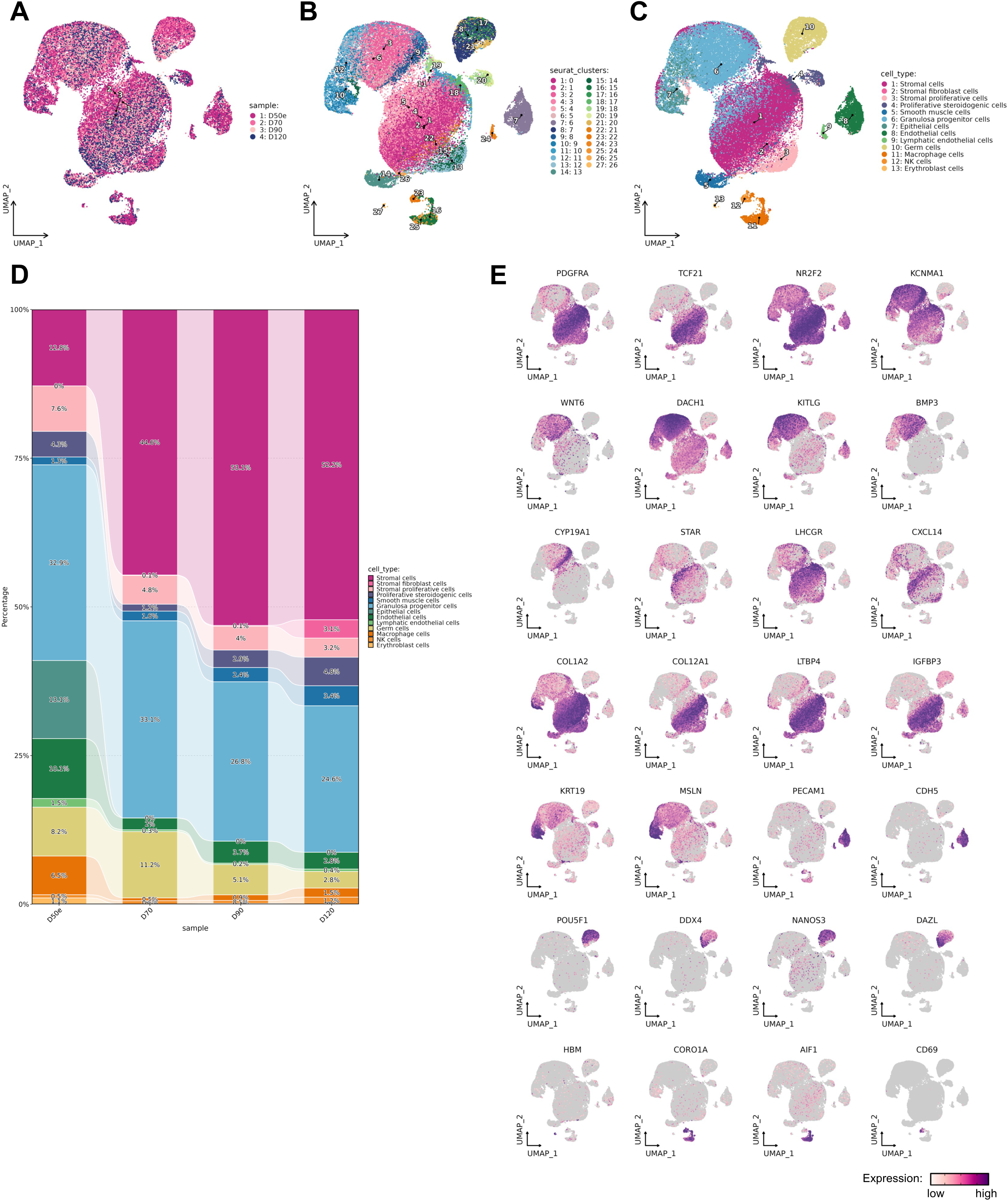
Single-cell RNA-sequencing analysis of bovine ovaries from 50–120 days of fetal age. (A) UMAP split by samples, (B) UMAP split by clusters, (C) UMAP split by cell identity annotation of the integrated dataset of bovine ovaries at day 50 (estimated; E), 70, 90, and 120 of fetal development. Each dot represents a single cell. Legend indicates cluster identities. (D) Proportion of cells in each cluster by sample age. Colors correspond to the age identified in the legend. (E) Feature plots showing relative expression of cells for markers of stromal cells (*PDGFRA, TCF21*), bipotential precursor cells (*NR2F2, KCNMA1*), granulosa progenitor cells (*WNT6, DACH1, KITLG, BMP3, CYP19A1, STAR, LHCGR, CXCL14*), stromal fibroblast cells (*COL1A2, COL12A1, LTBP4, IGFBP3*), epithelial cells (*KRT19, MSLN*), endothelial cells (*PECAM1, CDH5*), germ cells (*POU5F1, DDX4, NANOS3, DAZL*), erythroblasts (*HBM*), macrophages (*CORO1A, AIF1*), and NK cells (*CD69*).

### Distinct stages of germline development captured in the bovine fetal ovary

A total of 3,140 germ cells were captured by our analysis. By age, we could see fluctuations in this percentage, with germ cells comprising 8.2% (D50e), 11.2% (D70), 5.1% (D90), 2.8% (D120) of the ovary with increasing sample age from day 50-120. We surmise that this is reflective of the peak in mitotic activity from PGC arrival at the gonad until meiosis initiates around day 75, followed by increased apoptotic activity as only a fraction of oogonia give rise to oocytes within follicles, as well as growth of the ovary as a whole^36,37^. After filtering and sub-clustering, the germ cell population across all four timepoints resolved six transcriptionally distinct developmental stages (Figure 2A–C): PGCs (sub-clusters 0, 4, 5, 6, 7, 8, 9, 11, 14, and 15), transitioning oogonia (sub-clusters 1 and 13), proliferative oogonia (sub-cluster 2), committed oogonia (sub-clusters 3 and 10), meiotic prophase I oogonia (sub-cluster 12), and oocytes (sub-cluster 16). The temporal composition of each subtype shifted progressively across gestational ages (Figure 2D), with PGCs predominating at D50e and D70, and later-stage cell types (meiotic prophase I oogonia and oocytes) accumulating from D90 onward, consistent with the established timeline of bovine germline development^14,15,23,38^. PGCs expressed canonical markers, including *POU5F1, NANOG, SOX17, TFAP2C, PRDM1*, and *NANOS3,* together with the histone-related genes *H3C2* and *H3C8* and the transcriptional repressor *MXD3* (Figure 2E). These chromatin-associated markers are consistent with the broad epigenetic reprogramming characteristic of the PGC state. Transitioning oogonia showed elevated *PROM1, ALDH1L2, PSAT1,* and *IGFBP6*, indicating a metabolic shift and downregulation of early germline identity markers. Proliferative oogonia were characterized by *DPP10, HS3ST4*, and *CTNNA2*, consistent with active mitotic cycling and remodeling of cell adhesion. Committed oogonia, representing the transitional state to meiotic competence, showed strong upregulation of *DMRT1* and *DAZL* (Figure 2E, 2G). The appearance of *DMRT1* and *DAZL* expression in this subtype recapitulates findings in the human germline atlas, confirming conserved molecular checkpoints of oogonial commitment between cattle and primate species^39^.

**Figure 2.**
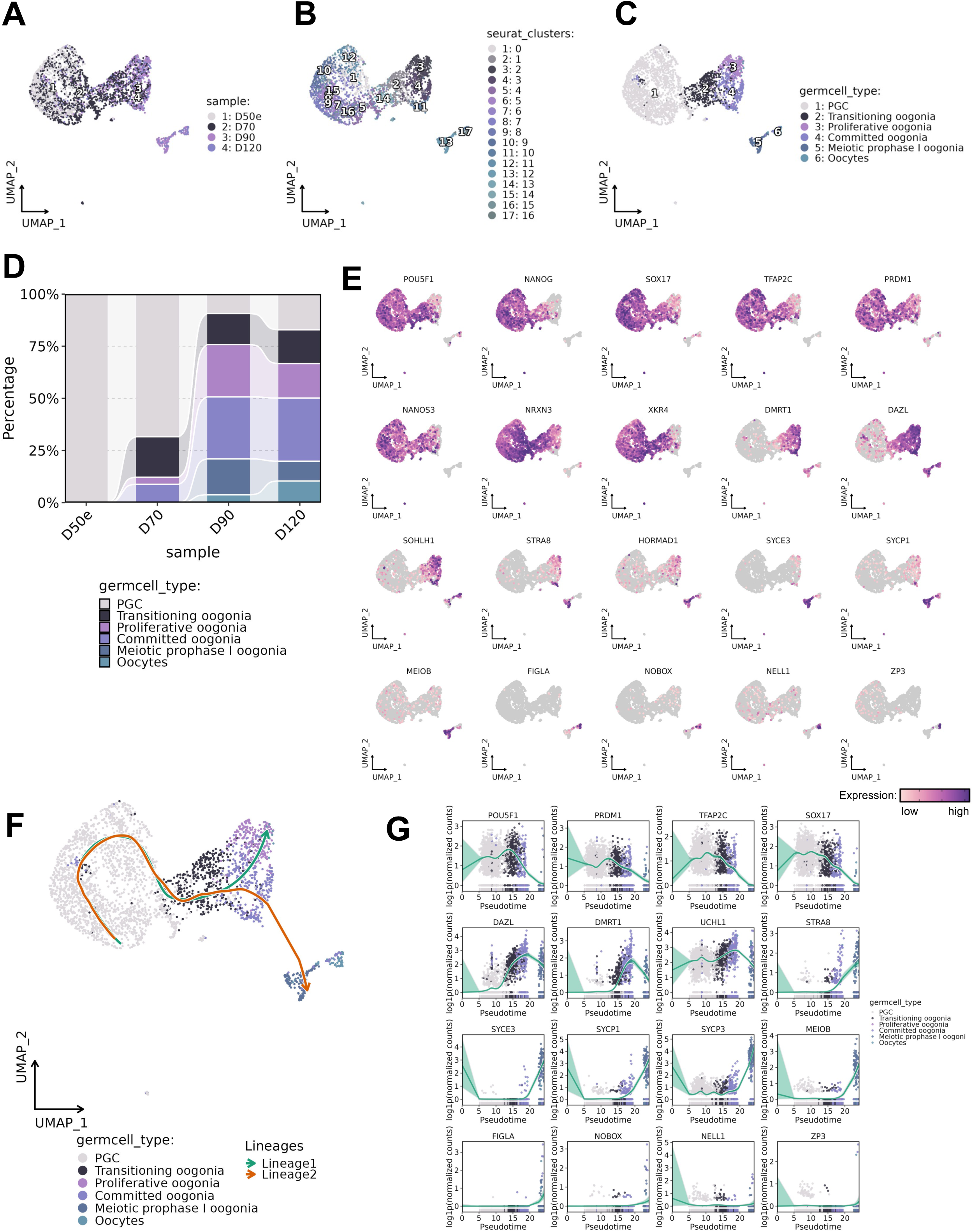
Single-cell RNA-sequencing analysis of bovine germ cell sub-cluster from 50–120 days of fetal age. (A) UMAP split by samples, (B) UMAP split by clusters, (C) UMAP split by cell identity annotation. Each dot represents a single cell. Legend indicates cluster identities. (D) Proportion of cells in each cell identity annotation cluster by sample age. Colors correspond to the age identified in the legend. (E) Feature plots showing relative expression of cells for markers of PGCs (*POU5F1, NANOG, NANOS3, TFAP2C, PRDM1, SOX17, NRXN3, XKR4*), proliferative and committed oogonia (*DMRT1, DAZL, SOHLH1*), meiotic oogonia (*STRA8, HORMAD1*, *SYCE3, SYCP1, MEIOB*), and oocytes (*FIGLA, NOBOX, NELL1, ZP3*). (F) Pseudotime trajectory of germ cells analyzed by Slingshot. (G) Pseudotime gene expression plots showing upregulation/downregulation of *POU5F1*, *PRDM1*, *TFAP2C*, *SOX17*, *DAZL*, *DMRT1*, *UCHL1*, *STRA8*, *SYCE3*, *SYCP1*, *SYCP3*, *MEIOB*, *FIGLA*, *NOBOX*, *NELL1*, *MOS* during development of PGCs to Oocytes.

Meiotic prophase I oogonia showed the highest fold-change differential expression of the entire dataset in key meiosis-specific genes, including *RAD51AP2, RAD21L1, SYCE3, SYCE1, MEIOB*, *SHOC1,* and *IHO1* (Figure 2E; Supplementary Figure 2A, 2C). These synaptonemal complex and meiosis I-specific transcripts confirm that the cells are in active prophase I of meiosis. The detection of these markers from D90 onward aligns with histological evidence of meiotic entry in bovine fetal ovaries between days 75 and 90^23,40^. The oocyte cluster showed the most divergent transcriptional signature in the entire germ cell trajectory and was defined by high expression of *FIGLA, NOBOX, ZP3*, and *NELL1*, in addition to reacquisition of *POU5F1* (Figure 2E). The oocyte-specific linker histone *H1-8* and nucleoplasmin *NPM2* point to active chromatin remodeling during oocyte formation. The transcription factors *FIGLA* and *NOBOX* are essential regulators of the oocyte-specific gene network and primordial follicle formation in mice and have been validated in human and non-human primates^41,42^. SCENIC regulon analysis (Supplementary Figure 2E) resolved a developmental progression that closely paralleled the transcriptional trajectory described above. A pluripotency/PGC-specifier module, comprising *POU5F1, NANOG, SOX15*, and *PRDM1*, showed activity restricted almost exclusively to the PGC population, consistent with the canonical primate PGC specification network and reinforcing the developmental identity assigned by marker-based clustering. A second module, including *ATF4*, *PBX1, ETS1*, and *FOXP1*, was broadly active across transitioning and proliferative oogonia, consistent with the proliferative, pre-commitment transcriptional state characteristic of these subtypes. A *HOX*-enriched module, comprising *HOXB7* and *HOXC9,* peaked specifically in committed oogonia, paralleling the HOX-family candidates (*HOXA5, HOXA10*, and *HOXB5*) independently identified by Pierson Smela et al. (2025) and further supporting *HOX* transcription factors as a conserved feature of oogonial commitment across species^41^. Finally, a distinct profile (*LEF1*, *FOXP2*, *ZBTB33, BCLAF1,* and *CHURC1)* marked meiotic prophase I oogonia, thematically consistent with the *HDAC*-associated chromatin remodeling factors flagged during meiotic entry in the same study^41^. Based on the small number of oocytes, we could not find any module for annotated oocytes (Supplementary Figure 2E). Together, these regulon-level findings corroborate with germ cell developmental stages and molecular transitions defined by differential expression and pseudotime analysis above. Trajectory analysis placed PGCs at the root; interestingly, two distinct pathways were observed. Lineage 1: with a continuous path through transitioning, towards the proliferative endpoint, and lineage 2: with a continuous path through transitioning, proliferative, and committed oogonia, diverging at the meiotic prophase I oogonia stage toward the oocyte endpoint (Figure 2F). Pseudotime-ordered heatmaps revealed distinct gene expression programs: an early wave of pluripotency-associated and epigenetic reprogramming genes (e.g., *POU5F1, NANOG, PRDM1*), a mid-trajectory wave of cell cycle and commitment genes (e.g., *DAZL, DMRT1, STRA8*), and a late-trajectory wave of meiotic and oocyte-specific programs (e.g., *SYCE3, FIGLA, ZP3*) (Figure 2G). Gene Ontology (GO) enrichment analysis of cluster-specific markers confirmed that the meiotic prophase I oogonia and oocyte populations were enriched for terms including “meiotic cell cycle,” “oogenesis,” “oocyte differentiation,” and “female gamete generation” (Supplementary Figure 2D). Additional differential gene expression and GO analyses for each germ cell subtype are presented in Supplementary Figure 3. Notably, PGCs were detected at all four gestational timepoints and still comprised a considerable fraction (17.0%) of the germ cell population at D120 (Supplementary Figure 2B).

Immunolocalization of “early” (POU5F1) and “late” (DAZL) stage germ cell proteins from estimated days 60–120 of bovine fetal ovarian development validated our sequencing findings that early germ cell states persist through day 120 of gestation and provided additional context that distinct germ cell states are spatially separated, with a gradient of PGCs (POU5F1+/DAZL-), transitioning oogonia (POU5F1+/DAZL+), committed oogonia (POU5F1-/DAZL+), and oocytes (POU5F1+/DAZL+, encircled by somatic support cells) from the outer–inner ovarian cortex (Supplementary Figure 3). As the ovary progresses between approximately D60 and D120 of fetal development, there is a clear shift from the predominance of POU5F1+/DAZL-cells (D60), appearance of POU5F1+/DAZL+ cells (D70), more prominent expression of DAZL and separation of DAZL+ and POU5F1+ cells (D90), and appearance of oocytes where POU5F1 expression returns (D90, D120), in line with our sequencing observations. This pronounced asynchrony of germ cell development, where PGCs persist alongside oocytes in the same tissue, is a distinctive feature of species with long gestation lengths and has been documented in humans, non-human primates, and cows^39,40,42^.

### Dynamics of emerging early supporting gonadal cells during bovine fetal ovarian development

After sub setting granulosa progenitor cells from the dataset, we re-clustered this population using the same pipeline and identified four transcriptionally-distinct subtypes (Figure 3A–C). We did not observe a major contribution from an individual sample across gestational ages (4,535 cells from D50e, 4,403 cells from D70, 3,330 cells from D90, and 3,343 cells from D120, representing 29.1%, 28.0%, 21.3%, and 21.4%, respectively, of the subset population) (Supplementary Figure 4B-C). The steroidogenic population, largely restricted to D50e (99.9%, Supplementary Figure 4B-C), showed high expression of *CYP17A1, DLK1, INSL3, ARX, LHCGR, STAR*, and *CYP11A1*. This signature points to an early, transient steroidogenic precursor, consistent with steroidogenic activity peaking around the time of sexual differentiation in the fetal ovary^43^. Pre-Granulosa 1 cells (PG1) emerged immediately following this steroidogenic wave, dominating the support-cell population at D70 (72.1%) before declining in relative proportion at D90 (18.3%) and D120 (15.8%). PG1 was defined by *KISS1, CDH4, RELN*, and *KLF5*. Its complete absence at D50e is consistent with the previous report indicating that support cells had not yet been captured at D50e^23^. Pre-Granulosa 2 cells (PG2) were minor at D50e (0.02%) and D70 (3.3%) but became the dominant subtype by D90 (56.5%) and D120 (65.8%) and were distinguished by the Notch target genes *HES4* and *HEY2* alongside the interferon-stimulated genes *IFI6, ISG15*, and *OAS2*, suggesting that active Notch-mediated intercellular communication accompanies this later, more committed state. Pre-Granulosa 3 cells (PG3) peaked transiently at D70 (19.6%) and D90 (20.9%) before declining sharply by D120 (8.2%), and expressed *CYP19A1* (aromatase), *MEIS2, GABRB1, PAPPA*, and *RSPO2*, pointing to active estrogen biosynthesis capacity and the most differentiated, follicle-associated identity among the pre-granulosa subtypes. Overall, the steroidogenic (29.1%), PG1 (27.6%), and PG2 (27.1%) populations were comparably represented across the full dataset, while PG3 (11.8%) and epithelial cells (4.5%) were comparatively minor fractions of the support-cell compartment. Granulosa cell development initiates after sexual differentiation and in preparation for and following the assembly of follicles^14^, which our dataset reflected in the sequential replacement of the steroidogenic population by PG1, then PG2, over the developmental time course. The epithelial cell subtype expressed *PODXL, KRT7, CDH6, KRT19*, and *UPK3B*, consistent with a surface epithelial identity distinct from the steroidogenic/granulosa lineage (Figure 3C–E). Additionally, the differential gene expression circle plot highlights the top 10 up/down regulated genes for each cell subtype (Figure 3F). Trajectory analysis of the pre-granulosa compartment placed steroidogenic population as the early-stage population, with PG2 and PG3 branching along a trajectory of increasing commitment (Figure 3G). Pseudotime-ordered gene expression dynamics revealed sequential activation of genes associated with follicle formation and gonadotropin responsiveness, including *FSHR, CYP19A1, PAPPA, MEIS2*, and *NR5A1*, at later pseudotime positions (Figure 3H). An additional heatmap of the DEG list and GO results is provided in Supplementary Figure 4. Interestingly, the identification of an epithelial cell subtype within the pre-granulosa compartment is consistent with recent findings in the human fetal ovary. Wamaitha et al. (2023) reported KRT19-positive epithelial cells as a spatially-defined component of the first-wave pre-granulosa population in human gonads from week 7 onward^39^. The presence of a *KRT19*-and *KRT7*-expressing epithelial subtype in the bovine fetal ovary at comparable developmental stages points to a conserved cell type within the somatic support niche of non-rodent species. Similarly, the bipotential somatic progenitor cells identified in the whole ovary dataset analysis by expression of *STAR, LHCGR*, and stromal markers are analogous to the early supporting gonadal cells (ESGCs) described by Wamaitha et al. (2023) and Garcia-Alonso et al. (2022) in humans, which emerge from stromal-identity progenitors and subsequently differentiate into organized pre-granulosa populations^39,44^. This parallel supports the bovine fetal ovary as a relevant model for studying the cell origins of the primate granulosa lineage.

**Figure 3.**
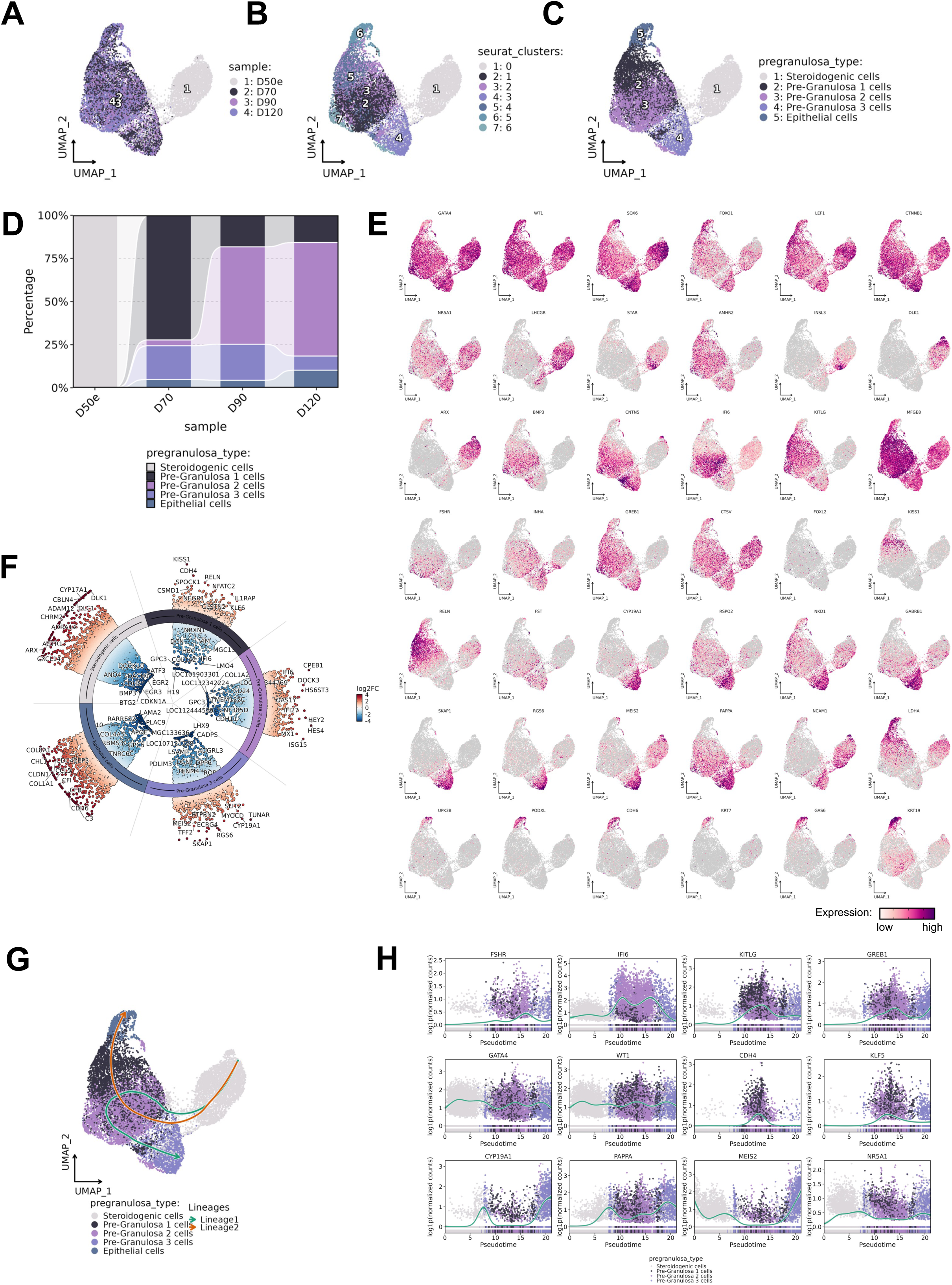
Single-cell RNA-sequencing analysis of early supporting gonadal cells (ESGCs) from day 50e0–120 of bovine fetal ovarian development. (A) UMAP split by samples, (B) UMAP split by clusters, (C) UMAP split by cell identity annotation. Each dot represents a single cell. Legend indicates cluster identities. (D) Proportion of cells in each cell identity annotation cluster by sample age. Colors correspond to the age identified in the legend. (E) Feature plots showing relative expression of cells for markers of steroidogenic cells (*STAR, CYP11A1, CYP17A1, INSL3, DLK1, ARX, LHCGR, NCAM1, CD24*), Pre-Granulosa 1 cells (*KISS1, CDH4, RELN, KLF5, ETV5*), Pre-Granulosa 2 cells (*HES4, HEY2, IFI6, FSHR, CPEB1*), Pre-Granulosa 3 cells (*CYP19A1, MEIS2, GABRB1, PAPPA, RSPO2*), and epithelial cells (*UPK3B, PODXL, CDH6, KRT7, KRT19*). (F) Volcano plot showing differential gene expression of individual identity annotation cluster versus others of sorted ESGCs. (G) Pseudotime trajectory of ESGCs analyzed by Slingshot. (H) Pseudotime gene expression plots showing upregulation/downregulation of *FSHR, IFI6, KITLG, GREB1, GATA4, WT1, CDH4, KLF5, CYP19A1, PAPPA, MEIS2*, and *NR5A1* during the development of ESGCs from day 50–120 of bovine fetal ovaries.

### Insights into cell–cell interactions between the forming germline and somatic niche of the ovary throughout key developmental hallmarks

To investigate the intercellular signaling environment of the developing bovine fetal ovary, we performed CellChat analysis on the full integrated dataset separated by gestational age. The total number of inferred interactions varied across D50e–D120 (2960, 2443, 2847, and 3081, respectively), reaching its lowest point at D70 and its highest at D120 (Figure 4A–B). Pathway-level comparison of relative information flow across all four timepoints revealed dynamic temporal regulation of distinct signaling programs (Figure 4C). BMP signaling was among the pathways active toward germ cells across this window, mediated through BMP5 at D50e and through both BMP5 and BMP2 at D120, acting via heteromeric BMPR1A/BMPR1B– ACVR2A/ACVR2B/BMPR2 receptor complexes on germ cells (Figure 4D–E). BMP signaling in the germline is known to regulate both PGC survival during the mitotic amplification phase and oogonia apoptosis, a required step in reducing germ cell number before follicle assembly^45^. KIT signaling showed a defined and consistent temporal pattern, with KITL (SCF)−KIT interactions detected as an incoming signal to germ cells at every timepoint from D50e through D120, consistent with the established role of this pathway in PGC survival and proliferation (Figure 4D–E). *KITL* expression by pre-granulosa cells at later timepoints suggests that, as the somatic niche matures, KIT signaling transitions from a broadly stromal-derived signal to a more granulosa-directed one, paralleling observations in human fetal ovarian development^39^. IGF signaling was present throughout the time course, though its receptor usage broadened over development: IGF2– IGF2R signaling was already detectable at D50e, while IGF1–IGF1R and IGF2–IGF1R interactions, along with the inhibitory IGFBP3–TMEM219, appeared from D70 onward and were sustained through D120 (Figure 4D–E). IGF signaling promotes germ cell proliferation and survival, and IGFBP3 is a high-affinity binding protein that modulates IGF bioavailability, consistent with a paracrine buffering mechanism for IGF activity within the developing niche. Midkine (MDK) signaling toward germ cells showed its broadest receptor repertoire at D50e, including MDK–ALK, MDK–PTPRZ1, MDK–SDC2, and MDK–SDC4 in addition to MDK– NCL and MDK–(ITGA6+ITGB1), before narrowing to MDK–NCL and MDK–(ITGA6+ITGB1) alone D70 through D120 (Figure 4D). MDK is a heparin-binding growth factor involved in cell migration, proliferation, and survival; the contraction of its receptor repertoire after D50e may reflect a narrowing of the paracrine signals available to germ cells as the niche matures. WNT signaling toward germ cells showed pathway-specific temporal restriction: WNT5A–FZD3 was detected at D50e, D90, and D120, while WNT6–(FZD3+LRP5/6) was restricted to the two mid timepoints (D70 and D90) and absent at both D50e and D120, consistent with a transient role for WNT6 in coordinating oogonial differentiation during the pre-meiotic window (Figure 4D). TENM–ADGRL2 signaling toward germ cells (TENM1, TENM2, TENM3, and TENM4 ligands from somatic populations acting on the latrophilin receptor ADGRL2 on germ cells) was present at D50e and sustained, with a slightly narrower ligand repertoire, through D70–D120. TENM– ADGRL interactions are implicated in cell adhesion and synaptic-like communication, and their presence in the ovarian niche across the full developmental window may reflect ongoing coordination of germ cell nest organization and cord formation.

**Figure 4.**
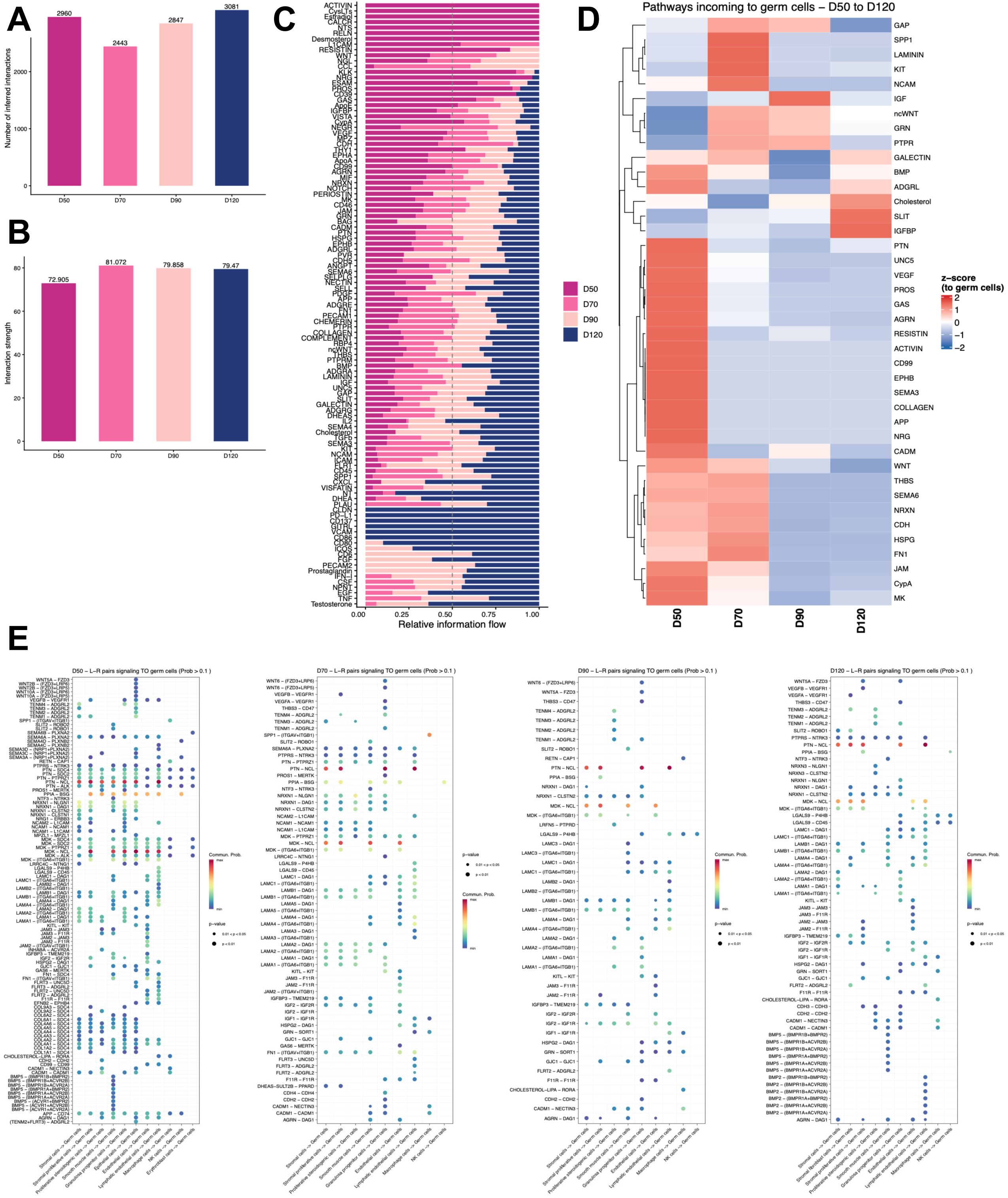
Cell–cell interactions between the forming germline and somatic niche of the bovine fetal ovary. (A) Barplot showing the total number of interactions per age. (B) Barplot showing interaction strength per age. (C) Barplot showing pathway-level contribution/comparison of all four time points. (D) Heatmap showing the relative signaling strength (z-score) of pathways incoming to germ cells across D50–D120. (E) Dot plots showing significant ligand–receptor pairs signaling to germ cells from each somatic population at D50, D70, D90, and D120; dot color indicates communication probability and dot size indicates significance.

## Discussion

This study presents a comprehensive single-cell transcriptomic atlas of the bovine fetal ovary, tracing the transitions of the germline from PGC colonization through primordial follicle formation, integrating data across four timepoints spanning gestational days 50 to 120. The resolution of six transcriptionally-distinct germ cell stages: PGCs, transitioning oogonia, proliferative oogonia, committed oogonia, meiotic prophase I oogonia, and oocytes, represents a novel and detailed subdivision of the bovine germline program and corroborates the markers and approximate timing of emergence of these cell types proposed by prior studies^14,15,22,23,40^. Notably, the committed oogonia stage shows a transcriptionally-distinct population that is intermediate between proliferative oogonia and meiotic prophase I oogonia. This stage is defined by co-expression of *DMRT1* and *DAZL*, and its identification is biologically meaningful: DMRT1 is required for germ cell sex determination and is expressed in oogonia before meiotic entry in multiple species, while DAZL marks the transition from pluripotency-associated to germline-committed identity^46–48^. The meiotic prophase I oogonia cluster shows the sharpest transcriptional transition in the entire dataset, with log2-fold changes exceeding 6–8 for multiple meiosis-specific genes, including *RAD51AP2, CIBAR2*, *RAD21L1, SYCE3, MEIOB*, and *IHO1*. These genes encode components of the synaptonemal complex, double-strand break repair machinery, and meiotic cohesion, all of which are essential for chromosome synapsis and recombination during prophase I. The oocyte cluster at the end of the pseudotime trajectory was defined by *FIGLA, NOBOX,* and *ZP3* expression. FIGLA and NOBOX are master regulators of the oocyte gene expression program in mice and are required for zona pellucida formation and primordial follicle assembly^49,50^. Their co-expression in the bovine oocyte cluster confirms conservation of this regulatory node. Pierson Smela et al. (2025) recently demonstrated that a combination of five transcription factors, including FIGLA, LHX8 and SOHLH1, can direct iPSC differentiation into DDX4-positive oogonia-like cells^41^. The transcriptional landscape of the bovine oocyte cluster identified here provides an in vivo reference point for benchmarking such in vitro-derived cell types and identifying which aspects of the oocyte program remain to be recapitulated.

One of the most striking features of this dataset is the presence of PGCs at all four timepoints, including D120, when meiotic prophase I oogonia and oocytes are also present. This pronounced asynchrony means that within a single D120 ovary, germ cells spanning the full developmental spectrum coexist. This contrasts with the mouse, where germline development is largely synchronized once germ cells enter the gonad^11^. Wamaitha et al. (2023) reported an analogous asynchrony in the human fetal ovary, with PGCs detected alongside meiotic cells at weeks 10–16^39^. Wamaitha et al. (2025) further extended this observation to the rhesus macaque, where germ cell developmental stages overlap extensively during the period from weeks 8–19 (∼D60–140)^42^. The current dataset establishes that bovine germline asynchrony follows a similar trajectory to primates. In addition to having similar gestation length and relative timing of developmental hallmarks, prior studies have suggested the cow as a relevant model of human reproduction due to similar ovarian architecture and follicular dynamics^51,52^. This is particularly important because human fetal tissue is rarely accessible for research beyond week 12–14, whereas bovine fetal ovaries at equivalent developmental stages are readily available.

CellChat analysis revealed a temporally dynamic intercellular signaling landscape, with BMP, KIT, IGF/IGFBP, and MDK pathways providing the core somatic–germline communication between D50e–D120. These findings both corroborate and extend observations from earlier studies. BMP signaling from the somatic niche to germ cells, primarily through BMP2/5 and heteromeric type I/II receptor complexes, was maintained across all timepoints. This is consistent with the established role of BMP signaling in PGC survival and in promoting apoptosis of nurse cells during germ cell nest breakdown^45^. The sustained BMP4 ligand–receptor activity in bovine germ cells at D90–D120, when follicle assembly is underway, is consistent with a model in which somatic BMP4 signaling finetunes the balance between germ cell survival and programmed attrition during the transition from cysts to individual follicles.

KIT–KITL signaling was present from D50e and showed temporal continuity across all ages, though the cellular sources of KITL evolved over time. At D50e–D70, *KITL* was broadly expressed by stromal and proliferative somatic cells. By D90–D120, *KITL* expression became concentrated in granulosa progenitor cell subtypes, reflecting the progressive organization of the somatic niche around forming follicles. The KIT receptor is a well-established survival factor in germ cells: in mice, KIT-null germ cells die before reaching the gonad, and KITL from granulosa cells is required for early follicle activation^53^. The bovine data show that the transition from a broadly distributed KITL source to a granulosa-restricted one occurs during the D70–D90 window, coinciding with the onset of organized pre-granulosa cell differentiation.

The distinction of MDK signaling in the incoming germ cell panel with MDK–NCL, MDK–PTPRZ1, and MDK–integrin interactions highlights ongoing communication between epithelial cells and the forming germline. MDK has received limited attention in the context of germline development, but its expression in ovarian stromal cells suggests it may function as a paracrine survival signal in the early ovarian niche^54^. Additionally, MDK supplementation in porcine PGC-like cells showed a mitogenic effect mediated through increased cell proliferation, decreased apoptosis, and meiotic inhibition by decreasing expression of DAZL and SCP3^55^.

WNT pathway activity, specifically non-canonical WNT5A–FZD3 and canonical WNT6– (FZD3+LRP5/6), were detected as incoming signaling to germ cells at D70 and D90, when most germ cells are in the transitioning and proliferative oogonia stages. WNT signaling is required for PGC specification in mice^56^ and has been implicated in bovine fetal ovarian development through single-cell transcriptomics^23^. The stage-specific enrichment of WNT inputs during the transitioning and proliferative oogonia period, identified here at the single-cell level for the first time in cattle, raises the question of whether WNT acts as a signal for oogonial identity maintenance or for controlling the timing of meiotic entry, a question that targeted functional studies could address.

Sub-clustering of the pre-granulosa compartment revealed five cell types: steroidogenic cells, PG1, PG2, PG3, and epithelial cells. This organization parallels the multi-wave model of pre-granulosa cell specification described in humans and non-human primates. Wamaitha et al. (2023) identified ESGCs as the bipotential progenitor population that gives rise to first-wave pre-granulosa cells, marked by *TSPAN8*, *LGR5*, and *FOXL2*, and discriminated from the somatic interstitium by *KRT19* expression starting at week 7 in humans^39^. In the current dataset, somatic progenitor cells expressing *STAR, LHCGR*, and stromal markers, are likely the bovine equivalent of these human ESGCs: a transitional somatic population retaining both stromal and gonadal identities before committing to the granulosa lineage.

The PG2 population, marked by *HES4, HEY2, CPEB1*, and *FSHR*, becomes a predominant cluster at D90 (when the first primordial follicles appear in the inner cortex^14^) and is consistent with a first-wave pre-granulosa identity that is responsive to FSH signaling (via *FSHR*) and actively engaged in Notch-mediated communication (via *HES4/HEY2*). In the mouse, first-wave bipotential pre-granulosa cells give rise to activated medullary follicles during mini puberty, while second-wave epithelial pre-granulosa cells populate the cortical reserve^57^. The PG3 population, with high *CYP19A1* and *MEIS2* expression, likely represent granulosa cells already associated with forming or nascent follicles at D90–D120. Interestingly, the epithelial cell subtype within the pre-granulosa compartment, expressing *KRT7* and *PODXL*, matches the *KRT19*-positive epithelial pre-granulosa cells described by Wamaitha et al. in human ovaries^39^. In that study, *KRT19* expression was the spatial discriminator of first-wave pre-granulosa cells within the cords from week 7 onward, and *KRT19*+/*FOXL2*+ cells were the dominant pre-granulosa cell type by week 15 in the rhesus macaque^42^. The bovine equivalents identified here suggest that the epithelial-to-mesenchymal transition-like dynamics of granulosa cell specification are conserved across cattle and primates. The transcriptional program of bovine PG1 cells, marked by *ETV5* and *CDH4*, is also notable. ETV5 is an ETS transcription factor required for granulosa cell function in mice downstream of FSH-FSHR signaling^58^. *CDH4* (R-cadherin) is a cell adhesion molecule enriched in germ cells and their supporting cells, and its expression in PG1 may reflect tight intercellular contact with germ cells within the ovigerous cords.

The bovine fetal ovary occupies a practical and scientifically relevant niche in reproductive biology. Bovine fetal tissue is a byproduct of livestock-derived food production and is available at multiple developmental stages that are ethically inaccessible in humans or technically challenging to obtain in non-human primates. The transcriptional parallels with human ovarian development documented here, including ESGC-like progenitors, multi-wave pre-granulosa subtypes, germline asynchrony, and conserved meiotic entry gene programs, strengthen the case for using bovine material as a proxy to answer questions about primate ovarian formation. From the assisted reproduction perspective, the six-stage germ cell atlas provides a reference map of the transcriptional states that must be recapitulated in vitro. For example, Pierson Smela et al. (2025) demonstrated that transcription factor-driven conversion of human iPSCs can produce oogonia-like cells in four days, but the resulting cells do not fully recapitulate the in vivo oocyte transcriptome^41^. The bovine clusters spanning PGCs to oocytes described here provide a high-resolution benchmark for evaluating the fidelity of such in vitro models and informing strategies to improve them.

## Conclusion

We present a temporal, single-cell transcriptomic atlas of the bovine fetal ovary spanning day 50 to day 120 of gestation, resolving 13 major cell populations and providing the first detailed characterization of six germ cell developmental stages and four pre-granulosa subtypes in cattle. The germ cell trajectory from PGCs through oocytes is defined by sequential transcriptional programs. The pronounced developmental asynchrony of the bovine germline, with PGCs persisting alongside oocytes at D120, is an intrinsic feature shared with primates. The five pre-granulosa subtypes: steroidogenic, PG1, PG2, PG3, and epithelial cells represent a parallel to ESGC-to-pre-granulosa progression described in human and primate fetal ovaries^39,42^, establishing the cow as a tractable model for the cellular and molecular origins of the ovarian granulosa lineage.

The cell–cell communication analysis reveals temporally specific deployment of BMP, KIT, IGF/IGFBP, MDK, and WNT signaling pathways, providing a systematic map of the somatic niche signals that accompany each phase of germ cell development. Together, these data advance fundamental understanding of bovine oogenesis, establish the intersection with human fetal ovarian development, and provide a resource for the rational design of in vitro culture systems aimed at recapitulating germ cell/somatic nursing cell progression.

## Acknowledgements

The authors would like to thank Juliana Candelaria for proofreading the manuscript.

## Author contributions

Conceptualization: C.G., R.C.B., R.B.A., A.C.D.; Methodology: C.G., R.C.B., R.B.A., B.B., B.K., S.P., C.K.G., S.R., A.C.D.; Formal analysis: C.G., R.C.B., A.C.D.; Investigation: C.G., R.C.B., R.B.A., J.M.S.; Resources: M.P., C.G., B.B., A.C.D.; Data curation: C.G., R.C.B., A.C.D.; Writing - original draft: C.G., R.C.B.; Writing - review & editing: C.G., R.C.B., R.B.A., J.M.S., S.R., B.K., S.P., B.B., C.K.G., S.R., A.C.D.; Visualization: C.G., R.C.B.; A.C.D.; Supervision: A.C.D.; Project administration: C.G., R.C.B.; A.C.D.; Funding acquisition: C.G., A.C.D.

## Data and resource availability

The raw and processed single-cell RNA-sequencing data generated in this study has been deposited in the NCBI Gene Expression Omnibus (GEO) database under accession number GSE344567. Any additional information will be available from the lead contact upon request.

## Competing interests

The authors have no competing interests to declare.

## Funding

This work was funded by Genus pIc. and USDA Hatch/Multistate project W-4171. C.G. was partially supported by the AFRI Predoctoral Fellowship (project award no. 2022-11304) from the U.S. Department of Agriculture, National Institute of Food and Agriculture.

---

**Supplementary Figure 1.** (A) UMAP of the integrated bovine ovary dataset showing sample distribution, Seurat clusters, and cell identity annotation. (B) Heatmap showing the top 15 gene markers identified in each bovine ovarian lineage during days 50–120 of development. (C) Bar plots showing specifically enriched GO pathways in each bovine ovarian lineage during days 50– 120 of development

**Supplementary Figure 2.** (A) Heatmap showing the top 40 gene markers identified in each germ cell sub-lineage during days 50–120 of development. (B) Number and percentage of germ cells per sub-lineage by sample age. (C) Volcano plot showing differential gene expression of individual identity annotation cluster versus others of sorted germ cells. (D) Bar plots showing specifically enriched GO pathways in each bovine germ cell sub-lineage during days 50–120 of development. (E) Heatmap showing the top regulatory transcription factors by SCENIC analysis in each germ cell sub-lineage during days 50–120 of development.

**Supplementary Figure 3.** Immunofluorescence-based protein localization of POU5F1 (red) and DAZL (green) at (A) estimated day 60, (B) day 70, (C) day 90, and (D) day 120 of bovine fetal ovarian development. Cortex regions (C) are defined in whole-mount images. The outer cortex (OC), mid-region cortex (MC), and inner cortex (IC) are highlighted in zoomed images. Overlay images include DAPI (blue). PGCs are indicated by yellow arrows; oocytes are indicated by white arrows. Scale bars: 500 μm.

**Supplementary Figure 4.** (A) Heatmap showing the top gene markers identified in each pre-granulosa sub-lineage during days 50–120 of development. (B, C) Number and percentage of cells per pre-granulosa sub-lineage by sample age. (D) Dotplot showing specifically enriched GO pathways in each bovine pre-granulosa sub-lineage during days 50–120 of development.

## Notes

### Competing Interest Statement

The authors have declared no competing interest.

## References

1. Anderson, R., Copeland, T.K., Schöler, H., Heasman, J., and Wylie, C. (2000). The onset of germ cell migration in the mouse embryo. Mech Dev 91, 61–68. 10.1016/s0925-4773(99)00271-3.

2. Guiltinan, C., Botigelli, R.C., Arcanjo, R.B., Candelaria, J.I., Lanzon, L.F., Smith, J.M., Becerra-Cortes, G., and Denicol, A.C. (2025). Molecular profiling of bovine primordial germ cell specification and migration onset reveals a conserved program in bilaminar disc embryos. Preprint at Developmental Biology, 10.1101/2025.06.04.657951 10.1101/2025.06.04.657951.

3. Irie, N., Weinberger, L., Tang, W.W.C., Kobayashi, T., Viukov, S., Manor, Y.S., Dietmann, S., Hanna, J.H., and Surani, M.A. (2015). SOX17 Is a Critical Specifier of Human Primordial Germ Cell Fate. Cell 160, 253–268. 10.1016/j.cell.2014.12.013.

4. Kobayashi, T., Zhang, H., Tang, W.W.C., Irie, N., Withey, S., Klisch, D., Sybirna, A., Dietmann, S., Contreras, D.A., Webb, R., et al. (2017). Principles of early human development and germ cell program from conserved model systems. Nature 546, 416–420. 10.1038/nature22812.

5. Tam, P.P., and Snow, M.H. (1981). Proliferation and migration of primordial germ cells during compensatory growth in mouse embryos. J Embryol Exp Morphol 64, 133–147.

6. Farini, D., Scaldaferri, M.L., Iona, S., La Sala, G., and De Felici, M. (2005). Growth factors sustain primordial germ cell survival, proliferation and entering into meiosis in the absence of somatic cells. Dev Biol 285, 49–56. 10.1016/j.ydbio.2005.06.036.

7. Nicholls, P.K., Schorle, H., Naqvi, S., Hu, Y.-C., Fan, Y., Carmell, M.A., Dobrinski, I., Watson, A.L., Carlson, D.F., Fahrenkrug, S.C., et al. (2019). Mammalian germ cells are determined after PGC colonization of the nascent gonad. Proc Natl Acad Sci U S A 116, 25677–25687. 10.1073/pnas.1910733116.

8. Irie, N., Lee, S.-M., Lorenzi, V., Xu, H., Chen, J., Inoue, M., Kobayashi, T., Sancho-Serra, C., Drousioti, E., Dietmann, S., et al. (2023). DMRT1 regulates human germline commitment. Nat Cell Biol 25, 1439–1452. 10.1038/s41556-023-01224-7.

9. Cantú, A.V., Altshuler-Keylin, S., and Laird, D.J. (2016). Discrete somatic niches coordinate proliferation and migration of primordial germ cells via Wnt signaling. J Cell Biol 214, 215–229. 10.1083/jcb.201511061.

10. McLaren, A. (1991). Development of the mammalian gonad: the fate of the supporting cell lineage. Bioessays 13, 151–156. 10.1002/bies.950130402.

11. Pepling, M.E., and Spradling, A.C. (1998). Female mouse germ cells form synchronously dividing cysts. Development 125, 3323–3328. 10.1242/dev.125.17.3323.

12. Koubova, J., Menke, D.B., Zhou, Q., Capel, B., Griswold, M.D., and Page, D.C. (2006). Retinoic acid regulates sex-specific timing of meiotic initiation in mice. Proc Natl Acad Sci U S A 103, 2474–2479. 10.1073/pnas.0510813103.

13. Bowles, J., Knight, D., Smith, C., Wilhelm, D., Richman, J., Mamiya, S., Yashiro, K., Chawengsaksophak, K., Wilson, M.J., Rossant, J., et al. (2006). Retinoid signaling determines germ cell fate in mice. Science 312, 596–600. 10.1126/science.1125691.

14. Fortune, J.E., Yang, M.Y., and Muruvi, W. (2010). The earliest stages of follicular development: follicle formation and activation. Soc Reprod Fertil Suppl 67, 203–216. 10.7313/upo9781907284991.018.

15. Lavoir, M.C., Basrur, P.K., and Betteridge, K.J. (1994). Isolation and identification of germ cells from fetal bovine ovaries. Mol Reprod Dev 37, 413–424. 10.1002/mrd.1080370408.

16. Wrobel, K.H., and Süss, F. (1998). Identification and temporospatial distribution of bovine primordial germ cells prior to gonadal sexual differentiation. Anat Embryol (Berl) 197, 451–467. 10.1007/s004290050156.

17. Bartholomew, R.A., and Parks, J.E. (2007). Identification, localization, and sequencing of fetal bovine VASA homolog. Anim Reprod Sci 101, 241–251. 10.1016/j.anireprosci.2006.09.017.

18. Hummitzsch, K., Irving-Rodgers, H.F., Hatzirodos, N., Bonner, W., Sabatier, L., Reinhardt, D.P., Sado, Y., Ninomiya, Y., Wilhelm, D., and Rodgers, R.J. (2013). A new model of development of the mammalian ovary and follicles. PLoS One 8, e55578. 10.1371/journal.pone.0055578.

19. Ideta, A., Yamashita, S., Seki-Soma, M., Yamaguchi, R., Chiba, S., Komaki, H., Ito, T., Konishi, M., Aoyagi, Y., and Sendai, Y. (2016). Generation of exogenous germ cells in the ovaries of sterile NANOS3-null beef cattle. Sci Rep 6, 24983. 10.1038/srep24983.

20. Luo, H., Zhou, Y., Li, Y., and Li, Q. (2013). Splice variants and promoter methylation status of the Bovine Vasa Homology (Bvh) gene may be involved in bull spermatogenesis. BMC Genet 14, 58. 10.1186/1471-2156-14-58.

21. Pennetier, S., Uzbekova, S., Perreau, C., Papillier, P., Mermillod, P., and Dalbiès-Tran, R. (2004). Spatio-temporal expression of the germ cell marker genes MATER, ZAR1, GDF9, BMP15,andVASA in adult bovine tissues, oocytes, and preimplantation embryos. Biol Reprod 71, 1359–1366. 10.1095/biolreprod.104.030288.

22. Planells, B., Gómez-Redondo, I., Sánchez, J.M., McDonald, M., Cánovas, Á., Lonergan, P., and Gutiérrez-Adán, A. (2020). Gene expression profiles of bovine genital ridges during sex determination and early differentiation of the gonads†. Biol Reprod 102, 38–52. 10.1093/biolre/ioz170.

23. Soto, D.A., and Ross, P.J. (2021). Similarities between bovine and human germline development revealed by single-cell RNA sequencing. Reproduction 161, 239–253. 10.1530/REP-20-0313.

24. Hayashi, K., Ohta, H., Kurimoto, K., Aramaki, S., and Saitou, M. (2011). Reconstitution of the mouse germ cell specification pathway in culture by pluripotent stem cells. Cell 146, 519–532. 10.1016/j.cell.2011.06.052.

25. Hikabe, O., Hamazaki, N., Nagamatsu, G., Obata, Y., Hirao, Y., Hamada, N., Shimamoto, S., Imamura, T., Nakashima, K., Saitou, M., et al. (2016). Reconstitution in vitro of the entire cycle of the mouse female germ line. Nature 539, 299–303. 10.1038/nature20104.

26. Botigelli, R.C., Guiltinan, C., Arcanjo, R.B., and Denicol, A.C. (2023). In vitro gametogenesis from embryonic stem cells in livestock species: recent advances, opportunities, and challenges to overcome. Journal of Animal Science 101, skad137. 10.1093/jas/skad137.

27. Shirasawa, A., Hayashi, M., Shono, M., Ideta, A., Yoshino, T., and Hayashi, K. (2024). Efficient derivation of embryonic stem cells and primordial germ cell-like cells in cattle. J. Reprod. Dev. 70, 82–95. 10.1262/jrd.2023-087.

28. Goszczynski, D.E., Denicol, A.C., and Ross, P.J. (2019). Gametes from stem cells: Status and applications in animal reproduction. Reproduction in Domestic Animals 54, 22–31. 10.1111/rda.13503.

29. Goszczynski, D.E., Cheng, H., Demyda-Peyrás, S., Medrano, J.F., Wu, J., and Ross, P.J. (2019). In vitro breeding: Application of embryonic stem cells to animal production. Biology of Reproduction 100, 885–895. 10.1093/biolre/ioy256.

30. Saragusty, J., Diecke, S., Drukker, M., Durrant, B., Friedrich Ben-Nun, I., Galli, C., Göritz, F., Hayashi, K., Hermes, R., Holtze, S., et al. (2016). Rewinding the process of mammalian extinction. Zoo Biol 35, 280–292. 10.1002/zoo.21284.

31. Zheng, G.X.Y., Terry, J.M., Belgrader, P., Ryvkin, P., Bent, Z.W., Wilson, R., Ziraldo, S.B., Wheeler, T.D., McDermott, G.P., Zhu, J., et al. (2017). Massively parallel digital transcriptional profiling of single cells. Nat Commun 8, 14049. 10.1038/ncomms14049.

32. Butler, A., Hoffman, P., Smibert, P., Papalexi, E., and Satija, R. (2018). Integrating single-cell transcriptomic data across different conditions, technologies, and species. Nat Biotechnol 36, 411–420. 10.1038/nbt.4096.

33. Germain, P.-L., Lun, A., Garcia Meixide, C., Macnair, W., and Robinson, M.D. (2021). Doublet identification in single-cell sequencing data using scDblFinder. F1000Res 10, 979. 10.12688/f1000research.73600.2.

34. Street, K., Risso, D., Fletcher, R.B., Das, D., Ngai, J., Yosef, N., Purdom, E., and Dudoit, S. (2018). Slingshot: cell lineage and pseudotime inference for single-cell transcriptomics. BMC Genomics 19, 477. 10.1186/s12864-018-4772-0.

35. Jin, S., Guerrero-Juarez, C.F., Zhang, L., Chang, I., Ramos, R., Kuan, C.-H., Myung, P., Plikus, M.V., and Nie, Q. (2021). Inference and analysis of cell-cell communication using CellChat. Nat Commun 12, 1088. 10.1038/s41467-021-21246-9.

36. Erickson, B.H. (1966). DEVELOPMENT AND RADIO-RESPONSE OF THE PRENATAL BOVINE OVARY. Reproduction 11, 97–105. 10.1530/jrf.0.0110097.

37. Shen, J., Cunha, G.R., Sinclair, A., Cao, M., Isaacson, D., and Baskin, L. (2018). Macroscopic whole-mounts of the developing human fetal urogenital-genital tract: Indifferent stage to male and female differentiation. Differentiation 103, 5–13. 10.1016/j.diff.2018.08.003.

38. Arcanjo, R.B., Guiltinan, C., Botigelli, R.C., Smith, J.M., and Denicol, A.C. (2026). Dynamics of germ cell development in the bovine fetal ovary. Sci Rep. 10.1038/s41598-026-63775-7.

39. Wamaitha, S.E., Nie, X., Pandolfi, E.C., Wang, X., Yang, Y., Stukenborg, J.-B., Cairns, B.R., Guo, J., and Clark, A.T. (2023). Single-cell analysis of the developing human ovary defines distinct insights into ovarian somatic and germline progenitors. Developmental Cell 58, 2097–2111.e3. 10.1016/j.devcel.2023.07.014.

40. Arcanjo, R.B., Guiltinan, C., Botigelli, R.C., Smith, J.M., and Denicol, A.C. (2026). Dynamics of Germ Cell Development in the Bovine Fetal Ovary. Preprint at In Review, 10.21203/rs.3.rs-9173445/v1 10.21203/rs.3.rs-9173445/v1.

41. Pierson Smela, M., Kramme, C.C., Fortuna, P.R.J., Wolf, B., Goel, S., Adams, J., Ma, C., Velychko, S., Widocki, U., Srikar Kavirayuni, V., et al. (2025). Rapid human oogonia-like cell specification via transcription factor-directed differentiation. EMBO Rep. 10.1038/s44319-025-00371-2.

42. Wamaitha, S.E., Rojas, E.J., Monticolo, F., Hsu, F., Sosa, E., Mackie, A.M., Oyama, K., Custer, M., Murphy, M., Laird, D.J., et al. (2025). Defining the cell and molecular origins of the primate ovarian reserve. Preprint at Developmental Biology, 10.1101/2025.01.21.634052 10.1101/2025.01.21.634052.

43. Dominguez, M.M., Liptrap, R.M., and Basrur, P.K. (1988). Steroidogenesis in fetal bovine gonads. Can J Vet Res 52, 401–406.

44. Garcia-Alonso, L., Lorenzi, V., Mazzeo, C.I., Alves-Lopes, J.P., Roberts, K., Sancho-Serra, C., Engelbert, J., Marečková, M., Gruhn, W.H., Botting, R.A., et al. (2022). Single-cell roadmap of human gonadal development. Nature 607, 540–547. 10.1038/s41586-022-04918-4.

45. Nilsson, E., Parrott, J.A., and Skinner, M.K. (2001). Basic fibroblast growth factor induces primordial follicle development and initiates folliculogenesis. Molecular and Cellular Endocrinology 175, 123–130. 10.1016/S0303-7207(01)00391-4.

46. Chen, H.-H., Welling, M., Bloch, D.B., Muñoz, J., Mientjes, E., Chen, X., Tramp, C., Wu, J., Yabuuchi, A., Chou, Y.-F., et al. (2014). DAZL Limits Pluripotency, Differentiation, and Apoptosis in Developing Primordial Germ Cells. Stem Cell Reports 3, 892–904. 10.1016/j.stemcr.2014.09.003.

47. Krentz, A.D., Murphy, M.W., Kim, S., Cook, M.S., Capel, B., Zhu, R., Matin, A., Sarver, A.L., Parker, K.L., Griswold, M.D., et al. (2009). The DM domain protein DMRT1 is a dose-sensitive regulator of fetal germ cell proliferation and pluripotency. Proc. Natl. Acad. Sci. U.S.A. 106, 22323–22328. 10.1073/pnas.0905431106.

48. Lin, Y., Gill, M.E., Koubova, J., and Page, D.C. (2008). Germ cell-intrinsic and -extrinsic factors govern meiotic initiation in mouse embryos. Science 322, 1685–1687. 10.1126/science.1166340.

49. Rajkovic, A., Pangas, S.A., Ballow, D., Suzumori, N., and Matzuk, M.M. (2004). NOBOX Deficiency Disrupts Early Folliculogenesis and Oocyte-Specific Gene Expression. Science 305, 1157–1159. 10.1126/science.1099755.

50. Soyal, S.M., Amleh, A., and Dean, J. (2000). FIGα, a germ cell-specific transcription factor required for ovarian follicle formation. Development 127, 4645–4654. 10.1242/dev.127.21.4645.

51. Adams, G.P., and Pierson, R.A. (1995). Bovine model for study of ovarian follicular dynamics in humans. Theriogenology 43, 113–120. 10.1016/0093-691X(94)00015-M.

52. Campbell, B.K., Souza, C., Gong, J., Webb, R., Kendall, N., Marsters, P., Robinson, G., Mitchell, A., Telfer, E.E., and Baird, D.T. (2003). Domestic ruminants as models for the elucidation of the mechanisms controlling ovarian follicle development in humans. Reprod Suppl 61, 429–443.

53. Hutt, K.J., McLaughlin, E.A., and Holland, M.K. (2006). Kit ligand and c-Kit have diverse roles during mammalian oogenesis and folliculogenesis. Mol Hum Reprod 12, 61–69. 10.1093/molehr/gal010.

54. Ikeda, S., and Yamada, M. (2014). Midkine and cytoplasmic maturation of mammalian oocytes in the context of ovarian follicle physiology. Br J Pharmacol 171, 827–836. 10.1111/bph.12311.

55. Shen, W., Park, B.-W., Toms, D., and Li, J. (2012). Midkine promotes proliferation of primordial germ cells by inhibiting the expression of the deleted in azoospermia-like gene. Endocrinology 153, 3482–3492. 10.1210/en.2011-1456.

56. Saitou, M., and Yamaji, M. (2010). Germ cell specification in mice: signaling, transcription regulation, and epigenetic consequences. Reproduction 139, 931–942. 10.1530/REP-10-0043.

57. Niu, W., and Spradling, A.C. (2020). Two distinct pathways of pregranulosa cell differentiation support follicle formation in the mouse ovary. Proc Natl Acad Sci U S A 117, 20015–20026. 10.1073/pnas.2005570117.

58. Akison, L.K., Alvino, E.R., Dunning, K.R., Robker, R.L., and Russell, D.L. (2012). Transient Invasive Migration in Mouse Cumulus Oocyte Complexes Induced at Ovulation by Luteinizing Hormone1. Biology of Reproduction 86. 10.1095/biolreprod.111.097345.

