## Supplementary figures and images for "Single-cell roadmap of bovine oogenesis and somatic niche interactions during fetal ovarian development"

### Supplemental Fig. 1

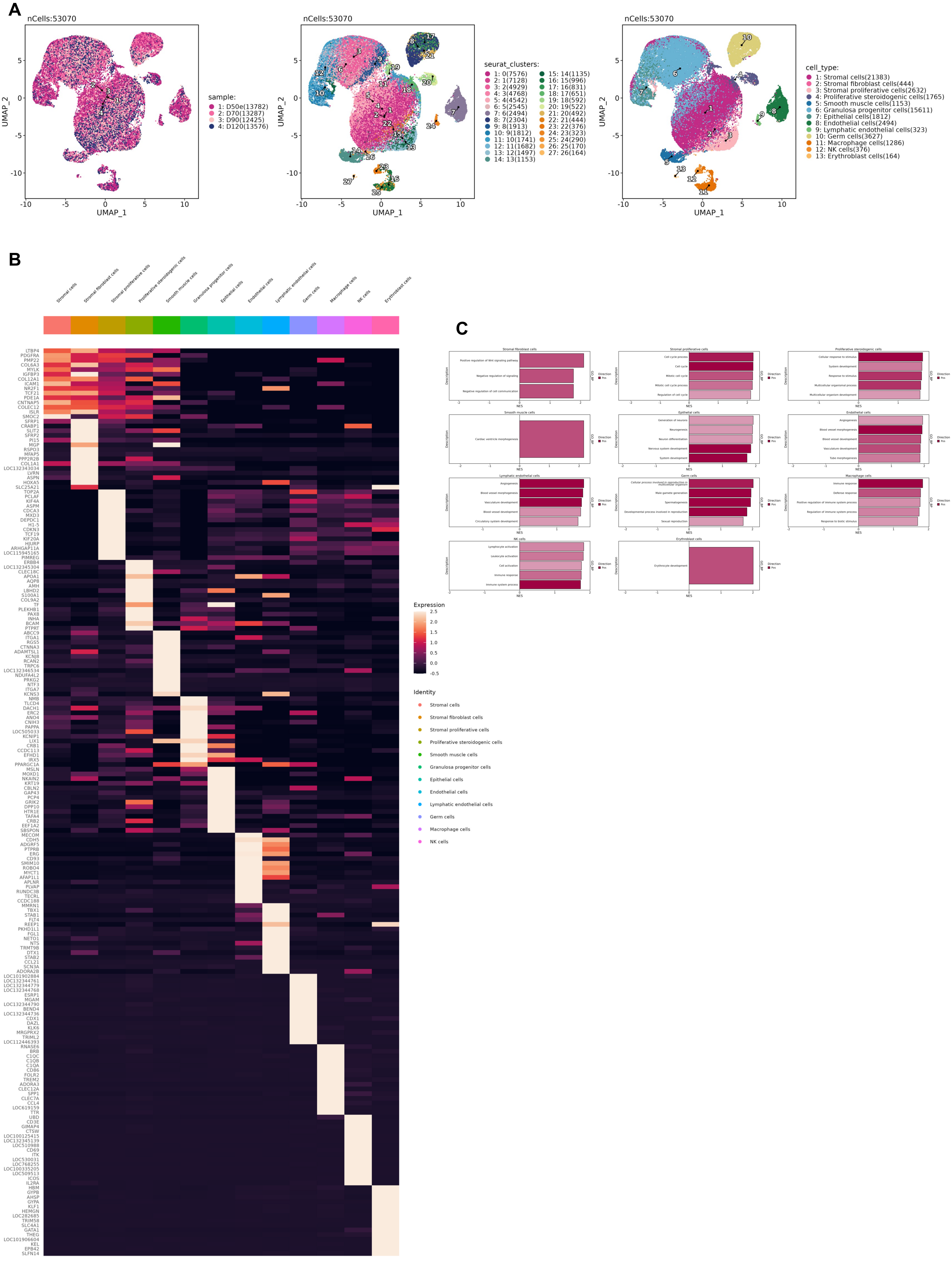

### Supplemental Fig. 3

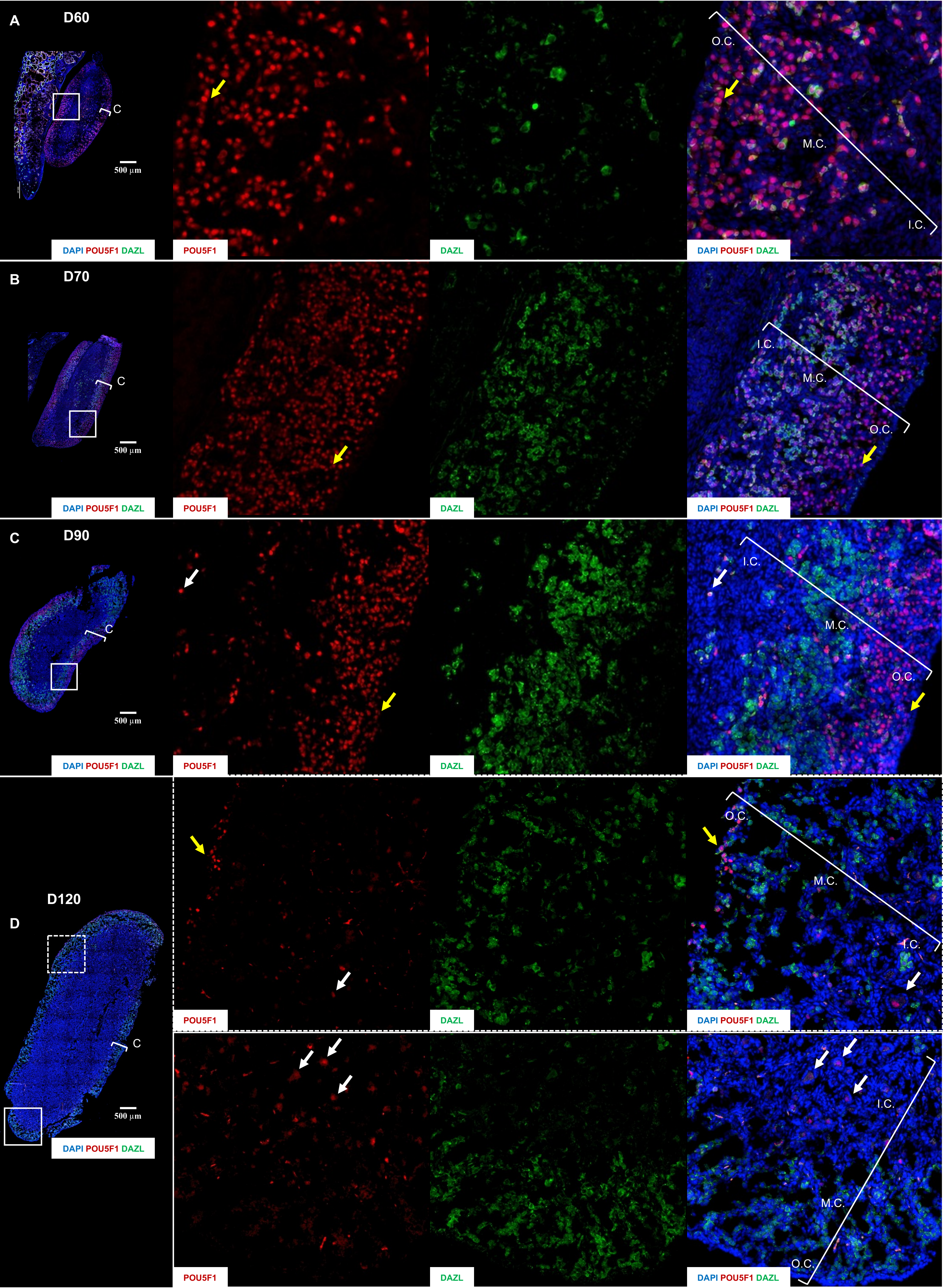
