## Supplemental Fig. 2 for "Single-cell roadmap of bovine oogenesis and somatic niche interactions during fetal ovarian development"

A

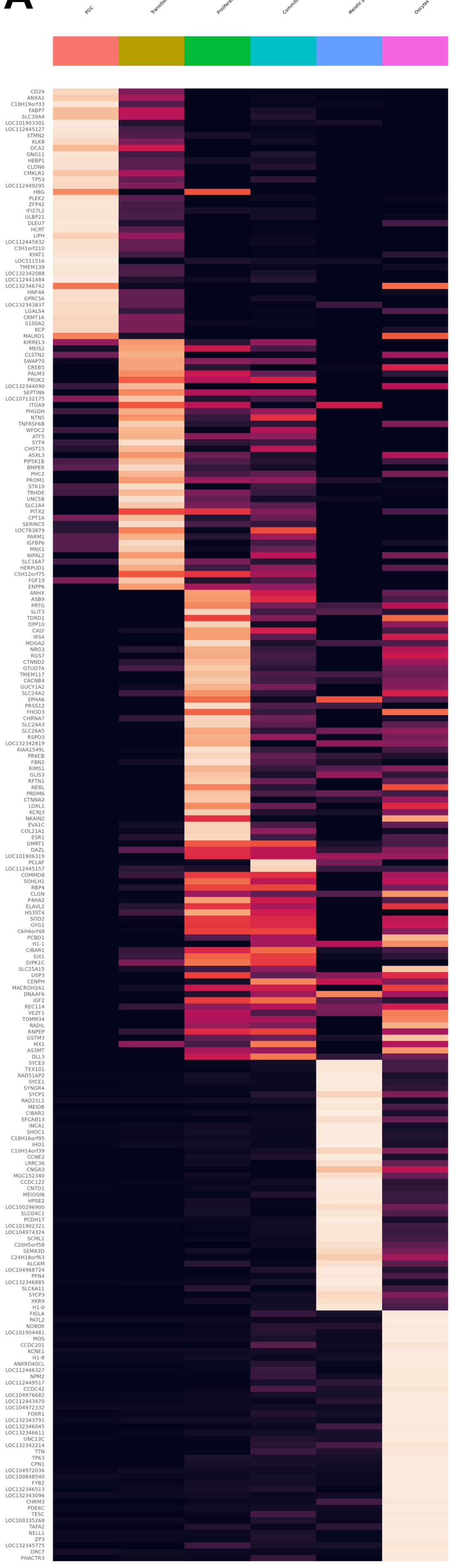


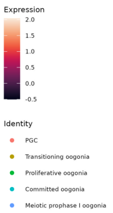


B

| sample | germcell_type | n | Percentage |
|---|---|---|---|
| D50e | PGC | 849 | 100 |
| D70 | PGC | 962 | 68.42 |
| D90 | PGC | 52 | 9.32 |
| D120 | PGC | 56 | 17.13 |
| D70 | Transitioning oogonia | 272 | 19.35 |
| D90 | Transitioning oogonia | 83 | 14.87 |
| D120 | Transitioning oogonia | 53 | 16.21 |
| D70 | Proliferative oogonia | 48 | 3.41 |
| D90 | Proliferative oogonia | 140 | 25.09 |
| D120 | Proliferative oogonia | 54 | 16.51 |
| D70 | Committed oogonia | 124 | 8.82 |
| D90 | Committed oogonia | 166 | 29.75 |
| D120 | Committed oogonia | 99 | 30.28 |
| D90 | Meiotic prophase I oogonia | 96 | 17.2 |
| D120 | Meiotic prophase I oogonia | 31 | 9.48 |
| D90 | Oocytes | 21 | 3.76 |
| D120 | Oocytes | 34 | 10.4 |

C

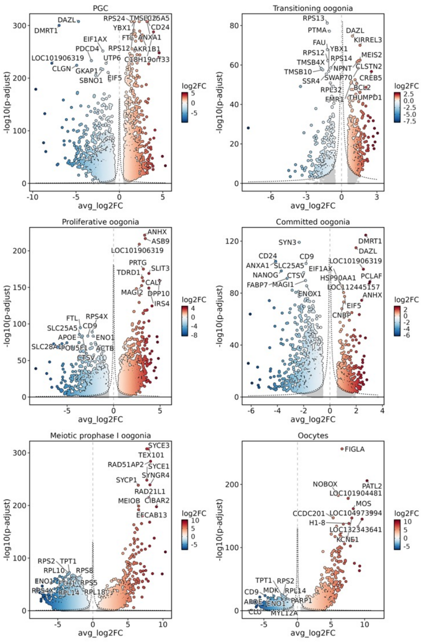


D

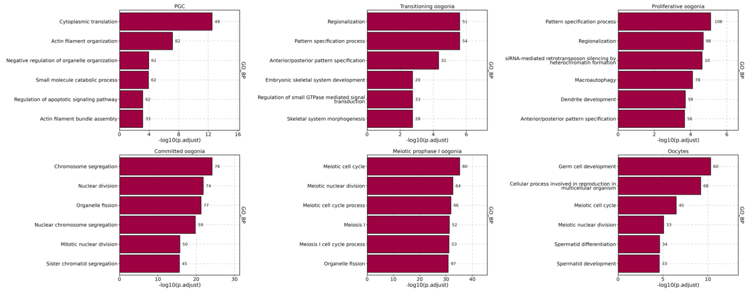


E

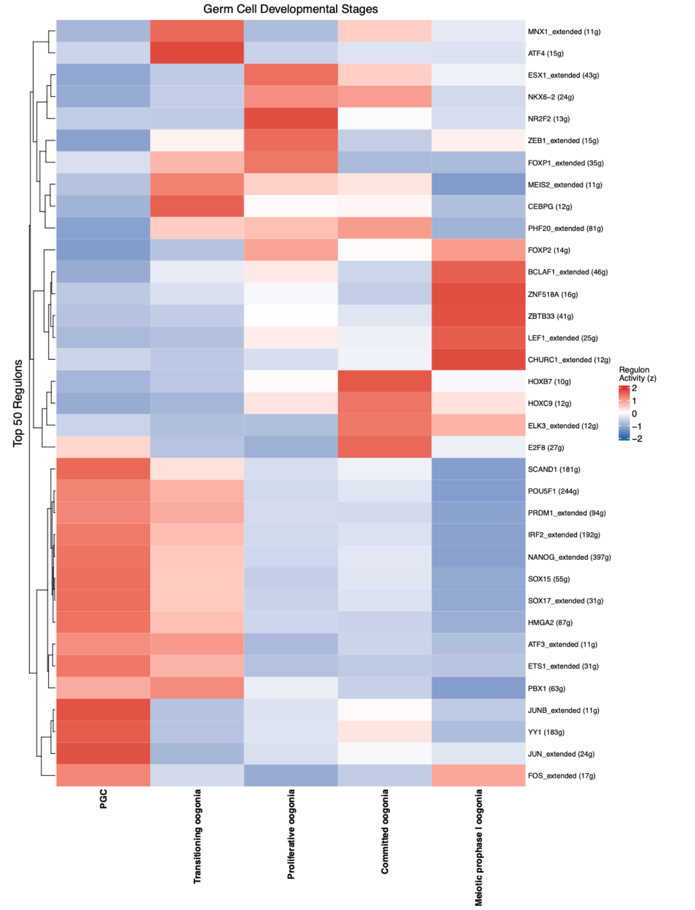
