## Supplemental Fig. 4 for "Single-cell roadmap of bovine oogenesis and somatic niche interactions during fetal ovarian development"

A

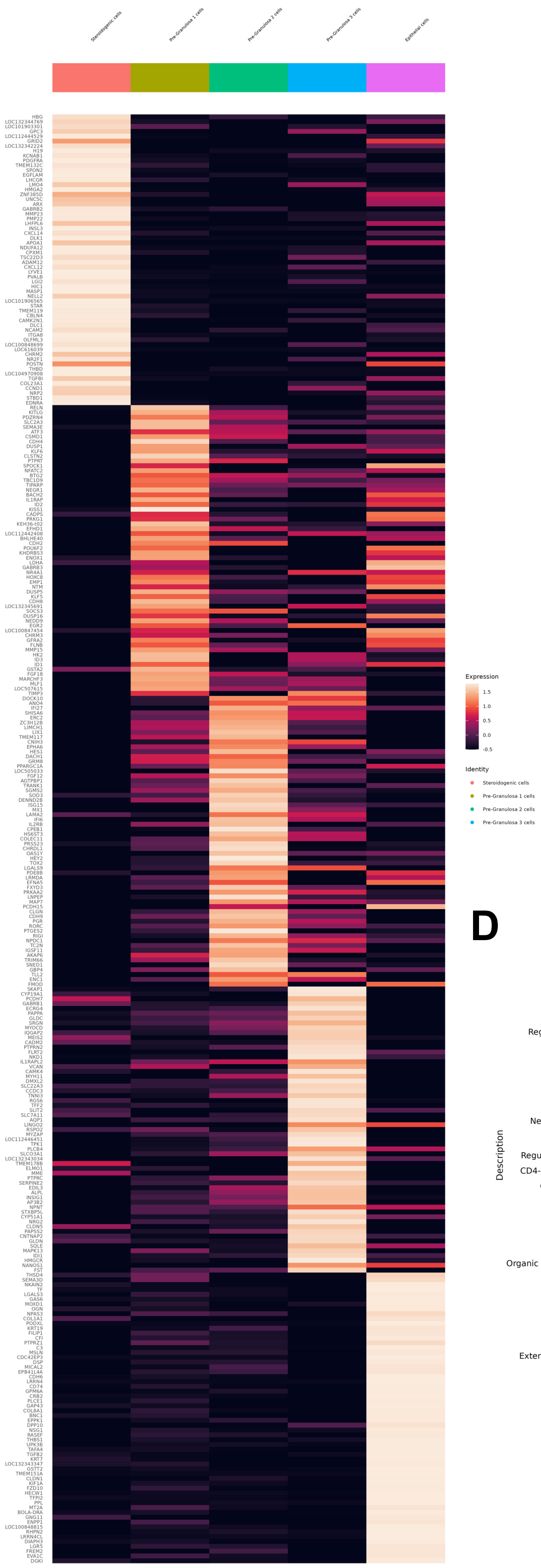


B

| sample | n | percentage |
|---|---|---|
| D50e | 4535 | 29.05 |
| D70 | 4403 | 28.2 |
| D90 | 3330 | 21.33 |
| D120 | 3343 | 21.41 |

C

| sample | pregranulosa_type | n | Percentage |
|---|---|---|---|
| D50e | Steroidogenic cells | 4532 | 99.93 |
| D70 | Steroidogenic cells | 9 | 0.2 |
| D90 | Steroidogenic cells | 1 | 0.03 |
| D70 | Pre-Granulosa 1 cells | 3173 | 72.06 |
| D90 | Pre-Granulosa 1 cells | 608 | 18.26 |
| D120 | Pre-Granulosa 1 cells | 528 | 15.79 |
| D50e | Pre-Granulosa 2 cells | 1 | 0.02 |
| D70 | Pre-Granulosa 2 cells | 147 | 3.34 |
| D90 | Pre-Granulosa 2 cells | 1881 | 56.49 |
| D120 | Pre-Granulosa 2 cells | 2200 | 65.81 |
| D50e | Pre-Granulosa 3 cells | 2 | 0.04 |
| D70 | Pre-Granulosa 3 cells | 862 | 19.58 |
| D90 | Pre-Granulosa 3 cells | 696 | 20.9 |
| D120 | Pre-Granulosa 3 cells | 275 | 8.23 |
| D70 | Epithelial cells | 212 | 4.81 |
| D90 | Epithelial cells | 144 | 4.32 |
| D120 | Epithelial cells | 340 | 10.17 |

D

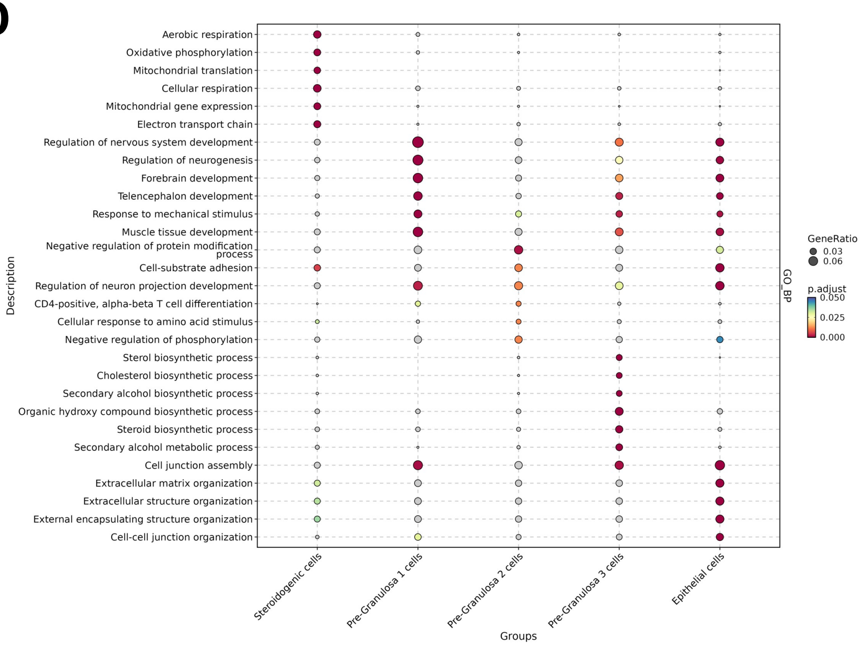
